# Phase-dependent suppression of beta oscillations in Parkinson’s disease patients

**DOI:** 10.1101/372599

**Authors:** Abbey B. Holt, Eszter Kormann, Alessandro Gulberti, Monika Pötter-Nerger, Colin G. McNamara, Hayriye Cagnan, Simon Little, Johannes A. Köppen, Carsten Buhmann, Manfred Westphal, Christian Gerloff, Andreas K. Engel, Peter Brown, Wolfgang Hamel, Christian K.E. Moll, Andrew Sharott

## Abstract

Synchronized oscillations within and between brain areas facilitate normal processing, but are often amplified in disease. A prominent example is the abnormally sustained beta-frequency (~20Hz) oscillations recorded from the cortex and subthalamic nucleus of Parkinson’s Disease patients. Computational modelling suggests that the amplitude of such oscillations could be modulated by applying stimulation at a specific phase. Such a strategy would allow selective targeting of the oscillation, with relatively little effect on other activity parameters. Here we demonstrate in awake, parkinsonian patients undergoing functional neurosurgery, that electrical stimulation arriving on consecutive cycles of a specific phase of the subthalamic oscillation can suppress its amplitude and coupling to cortex. Stimulus-evoked changes in spiking did not have a consistent time course, suggesting that the oscillation was modulated independently of net output. Phase-dependent stimulation could thus be a valuable strategy for treating brain diseases and probing the function of oscillations in the healthy brain.

## Introduction

Neural oscillations play a fundamental role in normal brain processing by temporally coordinating activity within and across regions (Buzsáki & Draguhn, 2004; Engel, Fries, & Singer, 2001). Dysfunctional communication resulting from an inability to properly modulate oscillatory activity, either through hypo- or hyper-synchrony, has been implicated in a number of neurological disorders (Schnitzler & Gross, 2005; Uhlhaas & Singer, 2006). Lesions, pharmacological treatments, and high frequency stimulation can all been used to disrupt irregular rhythmic activity, however these manipulations often result in wide-spread effects on network activity. Being able to selectively control synchrony without disruption to other physiological activity has the potential to improve therapies and insight into its role in normal functioning.

Functional neurosurgery for Parkinson’s Disease provides a unique opportunity to study the generation, propagation and perturbation of neuronal oscillations in the human brain. The implantation of deep brain stimulation (DBS) electrodes allows for both recording and electrical stimulation of basal ganglia nuclei. Such experiments have clearly demonstrated that the loss of midbrain dopamine neurons leads to abnormally sustained and synchronized beta oscillations (15-30 Hz) across the cortex and basal ganglia (Cassidy et al., 2002; Kuhn et al., 2005; Levy et al., 2002a; Sharott et al., 2018). These oscillations are thought to be mechanistically involved in symptom manifestation by distorting communication between brain areas needed for initiation of voluntary movement (Brown, 2007; Dorval & Grill, 2014; Engel & Fries, 2010).

The amplitude of beta oscillations correlates with severity of akinetic/rigid symptoms (Neumann et al., 2016; Sharott et al., 2014), and importantly, their reduction following high frequency (HF) (≥ 100 Hz) DBS positively correlates with motor improvement (Kuhn et al., 2008; Ray et al., 2008). While effective, HF DBS is limited by stimulation-induced side effects (Castrioto, Lhommee, Moro, & Krack, 2014; Hariz, Rehncrona, Quinn, Speelman, & Wensing, 2008; Tripoliti et al., 2011) and partial efficacy (Little & Brown, 2012). Triggering bursts of HF stimulation only during periods of high amplitude beta improves efficacy and reduces electrical energy delivered (Little et al., 2013); however, this could still disrupt physiological activity at timescales relevant for coding of movement in the subthalamic nucleus (STN) (Amirnovin, Williams, Cosgrove, & Eskandar, 2004; Lipski et al., 2017; Rodriguez-Oroz et al., 2005; Schrock, Ostrem, Turner, Shimamoto, & Starr, 2009; Sharott et al., 2018).

A phase-dependent approach, where stimulation is timed to a certain phase of the ongoing beta oscillation, has the potential to more selectively dampen the oscillatory activity. The utility of such a strategy can be seen in controlling tremor, where stimulation is locked to a specific phase of the behavioral oscillation (Cagnan et al., 2017). In many neurological disorders, such as akinesia and rigidity in PD, where no peripheral oscillation provides a marker of symptom severity, it may be necessary to time stimulation based on a neuronal oscillation (Azodi-Avval & Gharabaghi, 2015; Holt, Wilson, Shinn, Moehlis, & Netoff, 2016; Meidahl et al., 2017; Moll & Engel, 2017; Rosin et al., 2011). The approach is conceptually attractive, as it has the potential to modulate the timing of activity within and between structures, with less impact on gross excitability.

Using human intraoperative electrophysiological recordings, we demonstrate that in each parkinsonian patient there is a specific phase of the subthalamic LFP beta oscillation at which consecutive pulses of local electrical stimulation can suppress its amplitude without changes to overall excitability. Modulation extends to the output of STN neurons and to cortico-subthalamic synchrony. These results provide the first evidence in humans for using the phase of a subcortical oscillation to more selectively control its amplitude, and opens up the possibility of using such an approach for neurological disorders with oscillatory pathologies and to test the mechanistic role of these activities in functional processes.

## Materials and Methods

This study was conducted in agreement with the Code of Ethics of the World Medical Association (declaration of Helsinki, 1967) and was approved by the local ethics committee. All patients were previously diagnosed with advanced idiopathic Parkinson’s disease and gave their informed consent to participate. Recordings were made intra-operatively from ten patients undergoing awake surgery for bilateral implantation of DBS electrodes into the subthalamic nucleus (STN). Two patients were excluded from analysis for reasons discussed below.

### Patient information

Recordings were made while simultaneously delivering stimulation in ten patients (6 males, 4 females, average age: 62.1 years SD: 7.6 years). All patients had akinetic/rigid symptoms, had significant improvement of motor symptoms following levodopa intake (motor section (III) of the Unified Parkinson’s Disease Rating Scale), displayed no major cognitive decline (evaluated using the Mattis Dementia Rating Scale (Mattis, 1988)), and were awake during the surgical procedure. Participation in the study extended the surgical procedure by approximately 15-30 minutes. Every effort was made to keep additional time to a minimum, and stress level was continuously monitored using a verbally administered numerical rating scale to ensure any prolongation had no effect on the patient’s level of distress. Clinical details are summarized in Table 1.

### Surgical procedures

Stereotaxic bilateral implantation of DBS electrodes into the STN was performed under local anesthesia. Surgical procedures and targeting details have been previously described (Hamel et al., 2003; Moll et al., 2014). Briefly, prior to surgery patients stopped taking all anti-parkinsonian medication overnight. Surgical planning of the electrode trajectories was based on fused images of CT and MRI scans acquired the day of surgery. The stereotaxic targeting of STN was approximated based on the following coordinates: 11–13 mm lateral, 1–3 mm inferior, and 1–3 mm posterior to the midcommissural point. The trajectory was altered to avoid major blood vessels, sulci, and ventricles. Low-dose procedural sedation and analgesia with remifentanil was stopped prior to the microelectrode mapping procedure.

### Electrophysiological recording

Microelectrode recordings (Alpha Omega Neuroprobe, AlphaOmega, Nazareth, Israel) were performed along three parallel tracks arranged in a concentric array (Neuro Omega, Alpha Omega, Nazareth, Israel). The central electrode was aimed at the anatomically planned target and was separated by 2 mm from outer electrodes anteriorly in the parasagittal plane and laterally in the coronal plane. STN borders could be readily delineated based on elevated background activity levels (Moran, Bar-Gad, Bergman, & Israel, 2006) and characteristic firing properties of STN neurons (Sharott et al., 2014). Both unit activity and local field potentials (LFPs) were recorded from the microelectrode contact. Unit activity was bandpass filtered between 0.6 and 6 kHz, amplified (×20,000), and sampled at 44 kHz, while LFPs were bandpass filtered between 0.00070 and 0.4 kHz and sampled at 1.375 kHz. Recordings were referenced to the uninsulated distal most part of the guide tube for the corresponding microelectrode (macrotip diameter **~**0.8 mm, length ~ 1.5 mm, impedance <1 kΩ), located 3 mm above the microtip. EEG was recorded from scalp electrodes (needle electrodes) placed approximately at positions Fz, Cz, Pz, (according to the international 10-20 system), referenced to the nose. Signals were amplified (x 55,000), bandpass filtered between 0 and 0.3 kHz, and sampled at 1.375 kHz.

### Electrical stimulation of the dorsal STN area

Bipolar, biphasic, stimulation pulses were delivered through the macroelectrode contacts of two electrodes while the microelectrode recording contact of the third electrode was within the STN. Stimulation parameters were as follows: total pulse width: 200 μsec, 100 μsec initial phase negative,100 μsec positive phase; amplitude: 0.25 – 2 mA; constant current; stimulation time: 15-115 seconds, as permitted (all but one > 30 seconds). This resulted in stimulation being applied to the area immediately dorsal to the STN while LFPs and units were recorded from within the STN (Supplemental Fig. 1). Stimulation parameters were selected to modulate neuronal activity within the STN, while still allowing for a reliable LFP signal to be recovered from the recording electrode following stimulus artifact removal. Stimulation did not result in motor evoked potentials.

Stimulation was applied at or near the peak beta frequency (beta frequency stimulation) to determine effects of stimulation timing on beta oscillation amplitude. When well matched, stimulus pulses occurred at the same phase of the oscillation for at least two consecutive cycles. However, due to the natural variability in frequency and burst-like nature of the oscillation (Feingold, Gibson, DePasquale, & Graybiel, 2015; Tinkhauser et al., 2017), pulses drifted through different phases of the oscillation over the entire recording (Brittain, Probert-Smith, Aziz, & Brown, 2013; Cagnan et al., 2013). Only patients whose peak beta oscillation frequency was within 5 Hz of the stimulation frequency were included in analysis to ensure consecutive cycles of stimulation occurring at the same phase.

### Spike train processing

Spike (single and multi-unit) trains were separated from background activity using standard spike sorting procedures post-hoc (Spike2, Cambridge Electronic Design Limited) (Mallet, Pogosyan, Marton, et al., 2008; Mallet, Pogosyan, Sharott, et al., 2008), including template matching, principal component analysis, and supervised clustering. When a cluster was not separable, spike trains were defined as multi-units. Firing rates during stimulation were compared to rates before the onset of the first stimulus pulse at a given stimulus amplitude and depth. For the calculation of rates during stimulation, a 2.5 ms window following each pulse during which spikes could not be detected due to the resulting artifact was removed. The Wilcoxon’s signed rank test was used to evaluate effects of beta frequency stimulation on firing rate.

### Stimulus artifact removal

Data were analyzed offline using MATLAB (Mathworks, Natick, MA). A linear interpolation was used to remove sharp electrical artifacts in signals. To remove stimulus evoked artifacts seen in the LFP (Supplemental Fig. 2) a Kalman filter approach was used (Morbidi et al., 2007). The Kalman filter is a recursive approach which predicts the current state of the system and uses noisy measurements as feedback to update the prediction at each sample point. Briefly, we assume the recorded LFP is a summation of the unstimulated signal and the stimulus artifact. An autoregressive model was fit to a segment of unstimulated data and a transfer function model was fit to the average stimulus evoked artifact. The Kalman filter was then implemented and results used to estimate the artifact-free signal without phase distortion.

### Spectral power analysis

To evaluate overall effects of beta frequency stimulation on LFP beta power, spectra were normalized to the total power between 5 – 45 Hz and expressed as percentage of total power (%). Power between 0-5 Hz and above 45 Hz was eliminated to avoid contamination by movement and mains noise. The Wilcoxon’s signed rank test was used to evaluate statistical effects of stimulation on beta power.

### Instantaneous phase and amplitude estimation

To estimate the phase and amplitude of the beta oscillation, signals were bandpass filtered ±3 Hz around the peak beta frequency using a second order Butterworth filter with zero-phase digital filtering to preserve the true phase of the signal. The Hilbert transform was then used to estimate the instantaneous phase and envelope of the oscillation. Phase is defined as 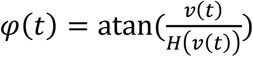, where *v*(*t*) is the filtered LFP signal and H(v(t))is the Hilbert transform of *v*(*t*). The amplitude envelope is defined as 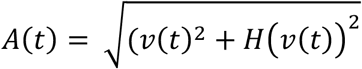.

### Instantaneous effects of stimulus phase

To assess how stimulus pulses occurring at a certain phase of the beta oscillation affect the amplitude envelope, stimulus phase was divided into eight overlapping phase bins ¼ of a cycle wide. The percentage change in median envelope over the cycle following the stimulus pulse was compared to the median envelope of the entire signal. Surrogate results were generated by sampling an unstimulated portion of data at the stimulation frequency. The Kruskal-Wallis test was use to assess phase dependent effects on beta amplitude in both the surrogate and stimulation conditions. Stimulation effects for each phase bin were compared to surrogates using the Wilcoxon rank-sum test.

### Cumulative effects of stimulus phase

To evaluate cumulative phase dependent effects of stimulation on beta amplitude, periods where stimulation occurred at the same phase (eight overlapping bins, ¼ of a cycle wide) coincidentally were used. As phase was not being tracked in real-time, the chances of observing further stimuli occurring within the same phase bin decreased as the number of stimuli increased (i.e. five consecutive stimuli in the same phase bin occurred less often than three). If there were fewer than five occurrences of one, two, three, four, or five consecutive pulses delivered at a specific phase throughout the entire recording, this occurrence was eliminated from analysis. For each patient, suppressing and amplifying bins were defined as the phase bins leading to the maximum suppression and amplification of the oscillation envelope in the LFP respectively. The Kruskal-Wallis test and multiple comparison test corrected using the false discovery rate were used to assess the significance of consecutive pulses at the amplifying and suppressing phase bin for each patient as well as across the group (Hochberg & Benjamini, 1990). Subsequently, to determine how precision of the defined stimulus phase affected beta amplitude modulation, the width of the amplifying and suppressing phase bins were widened and narrowed around the mean.

To ensure phase dependent modulation was not occurring in the spontaneous signal, cumulative effects were compared to surrogates. First, to verify that the number of consecutive stimuli seen in each bin does not simply lead to an increase or decrease in beta amplitude, the amplitude envelope during stimulation was replaced with an unstimulated segment from the same recording. Next, to ensure effects of stimulation were larger than any natural phase dependent variability in the oscillation amplitude, new amplifying and suppressing phase bins were identified using the unstimulated envelope sampled at the stimulation frequency (Supplemental Fig. 3). Stimulation effects were compared to surrogate effects using the Wilcoxon ranked sum test. Statistics reported in the results section are in comparison to the first surrogate.

To investigate whether phase changes accompany amplitude changes, we looked at the percentage of stimulus pulses that led to a phase slip when stimulating at either the suppressing or amplifying phase for consecutive cycles. Phase slips were identified when the instantaneous frequency (derivative of the unwrapped phase) exceeded two standard deviations above the mean, indicating a discontinuity in the oscillation phase (Pikovsky, Rosenblum, & Kurths, 2001). To attempt to avoid potential naturally occurring phase slips, only those occurring within 15 ms following the stimulus pulse were counted. Stimulus effects were compared to surrogates by replacing the instantaneous frequency with an unstimulated segment. Statistical effects were evaluated using the Wilcoxon ranked sum test.

To evaluate the effects of consecutive stimuli occurring at the defined amplifying and suppressing phase on local neuronal activity, background unit activity (BUA) was used. The BUA represents neuronal activity of a population of neurons around the recording contact, distinct from single and multiunit activity (Moran & Bar-Gad, 2010). To generate the BUA signal, large amplitude spikes (3 standard deviations above the mean) were removed from the unit microelectrode recording by replacing a window from 1 ms before to 3 ms after each spike with a random 4 ms spike-free segment of the same recording. The signal was then low-pass filtered at 300 Hz (third-order Butterworth, zero-phase digital filtering), rectified, and downsampled to 1.375 kHz (Moran & Bar-Gad, 2010; Moran, Bergman, Israel, & Bar-Gad, 2008; Sharott, Vinciati, Nakamura, & Magill, 2017). The BUA was bandpass filtered and analyzed for cumulative phase dependent effects of stimulation on signal amplitude as described for the LFP.

Midline EEG signals were used to assess cortico-subthalamic synchrony during stimulation at the defined amplifying and suppressing phase. EEG signals were band pass filtered between 0.001 kHz and 0.1 kHz to remove the contribution of slow drifts and high frequency activity, and notch filtered between 0.049 and 0.051 kHz, to remove line noise. To evaluate the phase relationship between the STN (LFP) and cortex (EEG), instantaneous phase of the EEG signal was determined as described for the LFP (2^nd^ order Butterworth bandpass filter, Hilbert transform). The phase synchrony index (PSI) between the two signals was then calculated over periods of three consecutive stimuli occurring at the suppressing or amplifying phase (Stam, Nolte, & Daffertshofer, 2007). The PSI is defined as:

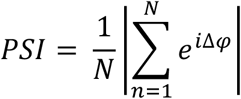

where N is the length of the segment (3 cycles), and ∆*φ* is the phase difference between the cortical and LFP signal (calculated using the Circular Statistics toolbox (Berens, 2009)). PSI values range from 0 to 1, with 1 representing a constant phase difference between the two signals.

## Results

Stimulation at or near the peak beta frequency was applied dorsal to the STN border in ten patients with Parkinson’s disease. When the stimulation frequency and ongoing oscillation were well matched, pulses could occur at a consistent phase for two or more cycles coincidentally, while drifting through all phases over the entire recording. Concurrently, STN local field potentials, STN unit activity, and mesial EEGs were recorded to investigate how stimulation timing interacts with the underlying neural beta oscillation (Fig. 1). Two patients were excluded from analysis; one did not have a significant beta oscillation (evaluated using the power spectrum of the STN LFP), and in the other the stimulus frequency was more than 5 Hz different from the peak oscillation frequency, preventing the stimulus phase from staying consistent within a quarter of a cycle over two or more consecutive beta cycles. In the eight patients included in analysis, the average oscillation frequency was 19 ± 5 Hz, and the stimulation frequency was 2.75 ± 1.75 Hz different from the peak beta frequency.

**Figure 1.**
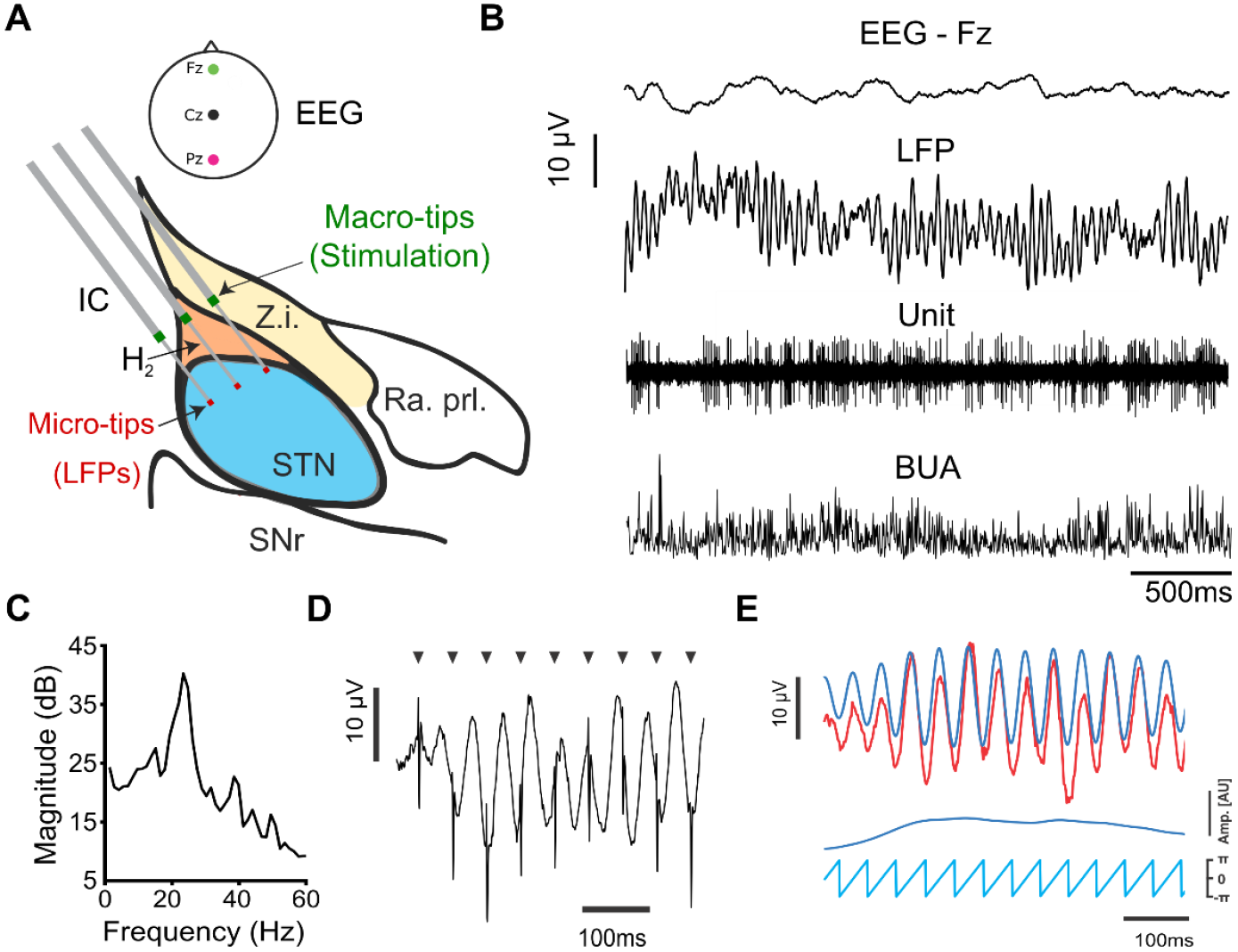
Cortico-subthalamic recordings during beta frequency stimulation in PD patients. (**A**) Schematic of the surgical setup shown on a sagittal view of the STN area, 11 mm lateral to the midline; modified from (Schaltenbrand and Bailey, 1959). Three microelectrodes were implanted along parallel tracks arranged in a concentric array. Stimulation was delivered through macro-tips located dorsal to the subthalamic nucleus while local field potentials were recorded from micro-tips within the subthalamic nucleus. Z.i.= zona incerta, IC = internal capsule, H_2_ = field H_2_ of Forel, Ra. Prl. = prelemniscal radiation, STN = subthalamic nucleus, SNr = substantia nigra pars reticulate. EEG was recorded from midline and frontal locations (Fz, Cz, and Pz). **(B)** Examples of signals recorded and used: EEG, local field potentials, unit activity, and background unit activity (BUA, generated using the unit channel). Example beta oscillation detected in the subthalamic local field potential recording. **(C)** A peak around 23 Hz can be seen in the power spectrum (corrected for 1/f falloff). **(D)** Bipolar, beta frequency stimulation (near peak beta frequency) was delivered through two macro-tips while local field potentials were recorded from the unstimulated micro-tip. **(E)** The signal (red) was bandpass filtered (blue) ± 3 Hz around the peak beta frequency. The Hilbert transform was used to estimate instantaneous phase (light blue) and amplitude (dark blue) from the filtered signal.

### Establishing stimulation parameters to investigate phase dependent effects

The overarching aim of this study was to examine whether the phase at which electrical stimulation was delivered relative to the underlying beta oscillation could produce short-latency effects on the amplitude of pathophysiological activity in the STN. To examine this question, we required a stimulation protocol that could modulate STN activity, but not cause gross changes over timescales of seconds. For example, other investigators have utilized stimulation protocols at beta frequency that reduce STN firing by up to 80% (Milosevic et al., 2018), which could confound the interpretation of changes in oscillatory activity. Similarly, STN stimulation can lead to large spectral peaks in the LFP, which would make analysis of stimulation effects on the ongoing oscillation difficult.

Following stimulus artifact removal, the spectral content of the recovered LFP was similar to that observed without stimulation (Fig. 2A). In line with this observation, stimulation did not result in a significant difference in either the peak beta power (± 3 Hz, p = 0.6665, Wilcoxon ranked sum test) (Fig. 2B) or wide band beta activity (8-35 Hz, p = 0.4894, Wilcoxon ranked sum test) (Fig. 2C), relative to total power between 5 and 45 Hz in seven out of the eight patients included in analysis. Additionally, beta frequency stimulation did not consistently alter the firing rate (Fig. 2D) of STN units (p = 0.9534, Wilcoxon ranked sum test) with respect to the unstimulated baseline.

**Figure 2.**
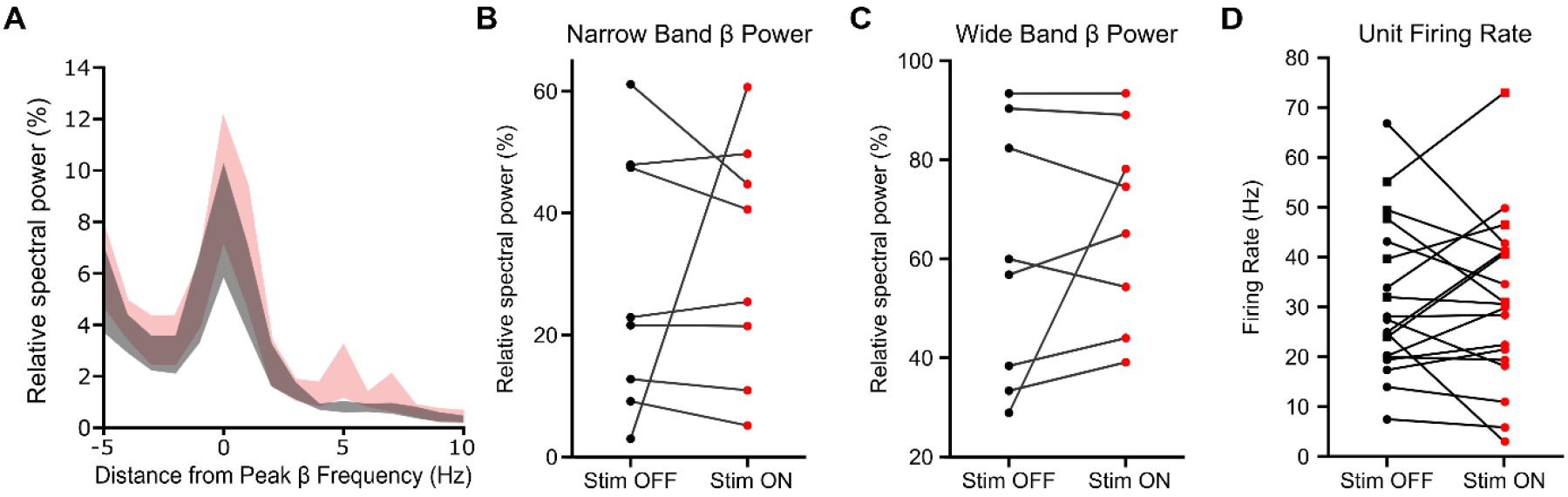
Beta frequency stimulation does not consistently modulate STN activity on the order of seconds. **(A)** Average ± S.E.M power spectra aligned to the peak beta frequency shows spectral activity calculated across the entire recording did not change significantly with beta frequency Stim ON across eight patients. A prominent beta peak was seen in Stim OFF (black) and Stim ON (red) conditions. **(B)** There was no significant difference in peak beta power (peak beta frequency ± 3 Hz) relative to 5-45 Hz with stimulation (red) (p = 0.6665, Wilcoxon ranked sum test). **(C)** There was no significant difference in total wide band beta frequency power (8-35 Hz) relative to 5-45 Hz with stimulation (p = 0.4894, Wilcoxon ranked sum test). **(D)** There was no significant difference in firing rates of putative subthalamic units between Stim OFF and Stim ON periods (p = 0.9534, Wilcoxon ranked sum test). Circles indicate cells classified as single units, squares multi-units.

While STN neurons were not modulated on the timescale of seconds, stimulation could lead to short-latency excitation (Fig. 3F) or inhibition of spiking (Fig. 3A, E, I), often followed by further multiphasic responses (Fig. 3C, G). Responses were in line with previous studies which show that STN neurons typically respond to afferent input with complex multiphasic responses resulting from interaction of monosynaptic and reciprocal polysnaptic input with their intrinsic pacemaker-driven firing (Magill, Sharott, Bevan, Brown, & Bolam, 2004; Nambu, Tokuno, & Takada, 2002; C. J. Wilson & Bevan, 2011). The magnitude of these responses could be amplitude dependent (Supplemental Fig. 4); however, there was no clear consistency based on stimulation/recording location (Supplemental Fig. 5). These results demonstrate that the stimulation protocol employed could alter the spike timing of individual STN neurons, without gross changes in firing rate or LFP beta power with respect to baseline.

**Figure 3.**
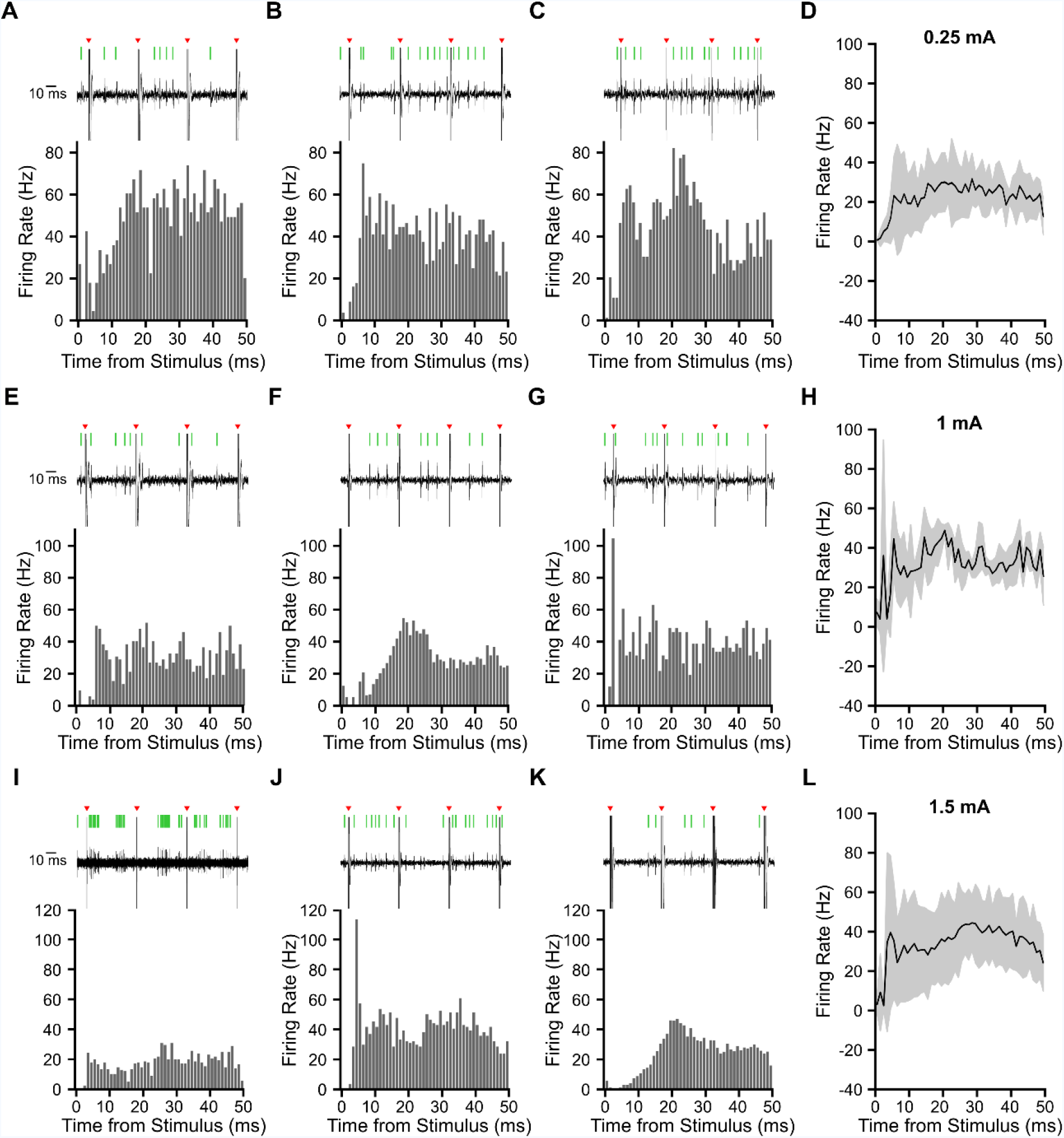
Beta frequency stimulation delivered dorsal to the subthalamic nucleus (STN) modulates STN unit activity. Peristimulus time histograms (PSTH), using 1 ms wide bins, from nine example STN units (single or multi-units) across seven patients. Beta frequency stimulation was applied at 0.25 mA **(A-C)**, 1 mA **(E-G)**, and 1.5 mA **(I-K)**. Spikes were detected from microelectrode recordings in the STN; representative examples of raw unit data during 3 consecutive electrical stimuli are shown above each PSTH (black: raw trace; red arrow: stimulation; green line: detected spike). **(D, H, L)** Average (± standard deviation) PSTH in response to 0.25 mA (7 units, 3 patients), 1 mA (3 units, 2 patients), and 1.5 mA (8 units, 4 patients) stimulation.

### Phase-dependent modulation of beta amplitude

Once optimal stimulus parameters were established, we investigated transient modulation of the STN LFP beta power by accounting for the phase relation between the stimulus pulse and ongoing oscillation. By using the phase of each independent stimulus pulse without accounting for the phase history of past stimuli, we found a significant trend in phase dependent modulation of the beta amplitude (Fig. 4) (p = 1.88e-4, Kruskal-Wallis test). However, modulatory effects were not significantly different from effects seen in an unstimulated portion of the recording sampled at the stimulation frequency (p > 0.05, Wilcoxon ranked sum test). This suggests there were no significant phase-dependent effects of single stimuli on the amplitude of the beta oscillation beyond what is seen in the natural variability of the signal.

**Figure 4.**
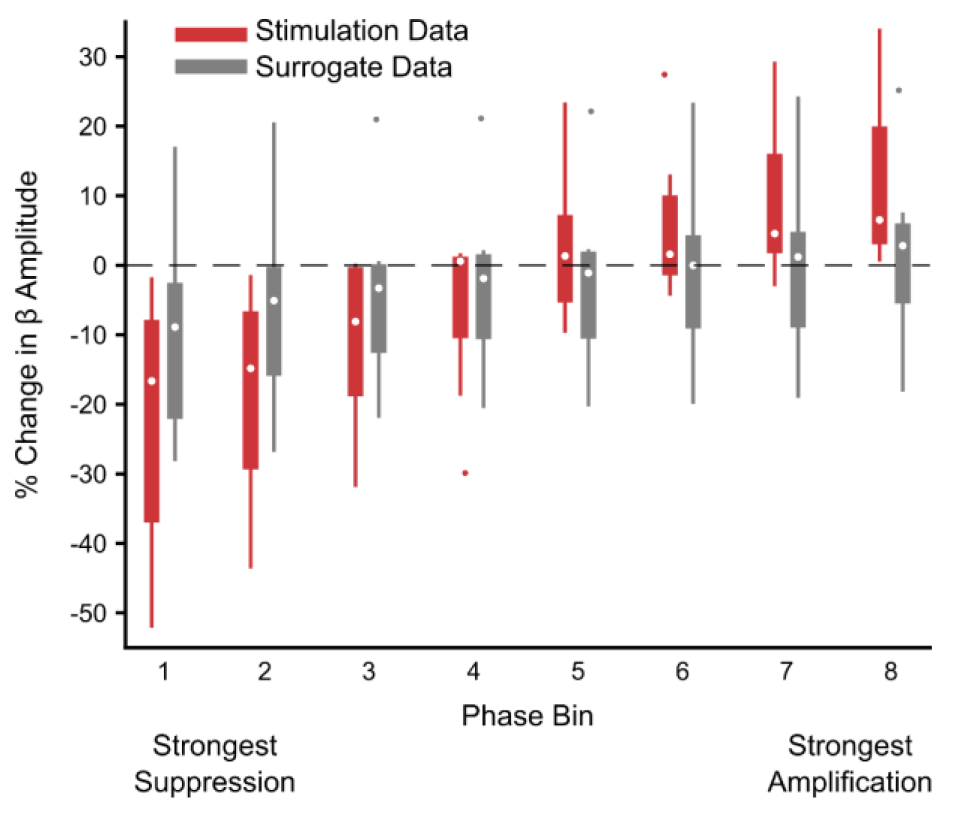
Phase dependent effects of single stimuli on beta amplitude do not exceed variability of the unstimulated LFP. To evaluate the effects of stimulus phase on beta amplitude, stimulus pulses across the entire recording were grouped into eight overlapping phase bins, ¼^th^ of a cycle wide. Across eight patients, bins were grouped according to median effect on beta amplitude, with bins leading to maximum suppression grouped together, etc. The percentage change in beta amplitude was dependent on stimulus phase (p = 1.88e-4, Kruskal-Wallis test) during stimulation, but not when sampling an unstimulated time-matched portion of the recording at the stimulation frequency (p = 0.3073, Kruskal-Wallis test). However, modulation from stimulation was not significantly different from that seen using the surrogate for any phase bin (p > 0.05, Wilcoxon ranked sum test). This indicates there were no significant phase dependent effects of stimulation on beta amplitude when stimulus phase history is not considered. Data is shown using a boxplot where the central dot is the median and box edges are the 25^th^ and 75^th^ percentiles. Outliers are plotted individually and defined as outside *q*_75_ – *w* * (*q*_75_ – *q*_25_) and *q*_25_ + *w \** (*q*_75_ – *q*_25_), where *q*_25_ and *q*_75_ are the 25^th^ and 75^th^ percentiles respectively and *w* is the maximum whisker length.

Because the stimulation frequency was well matched to the oscillation frequency, stimuli occurred at the same phase for consecutive beta cycles coincidentally. This allowed us to investigate modulatory effects during epochs when the stimulation phase was consistent within a quarter of a cycle for two or more consecutive beta cycles. In individual subjects, 2-3 consecutive pulses at a given phase could either suppress (suppressing phase) or amplify (amplifying phase) the beta amplitude of the following cycle compared to the median (Fig. 5A). Importantly, similar numbers of consecutive stimuli delivered at alternative phases did not result in a change in the amplitude. The phase bins leading to suppression and amplification of the beta oscillation were specific to each individual patient (Fig. 5C), but the difference between the two was always between 90 and 180 degrees (Fig. 5D).

**Figure 5.**
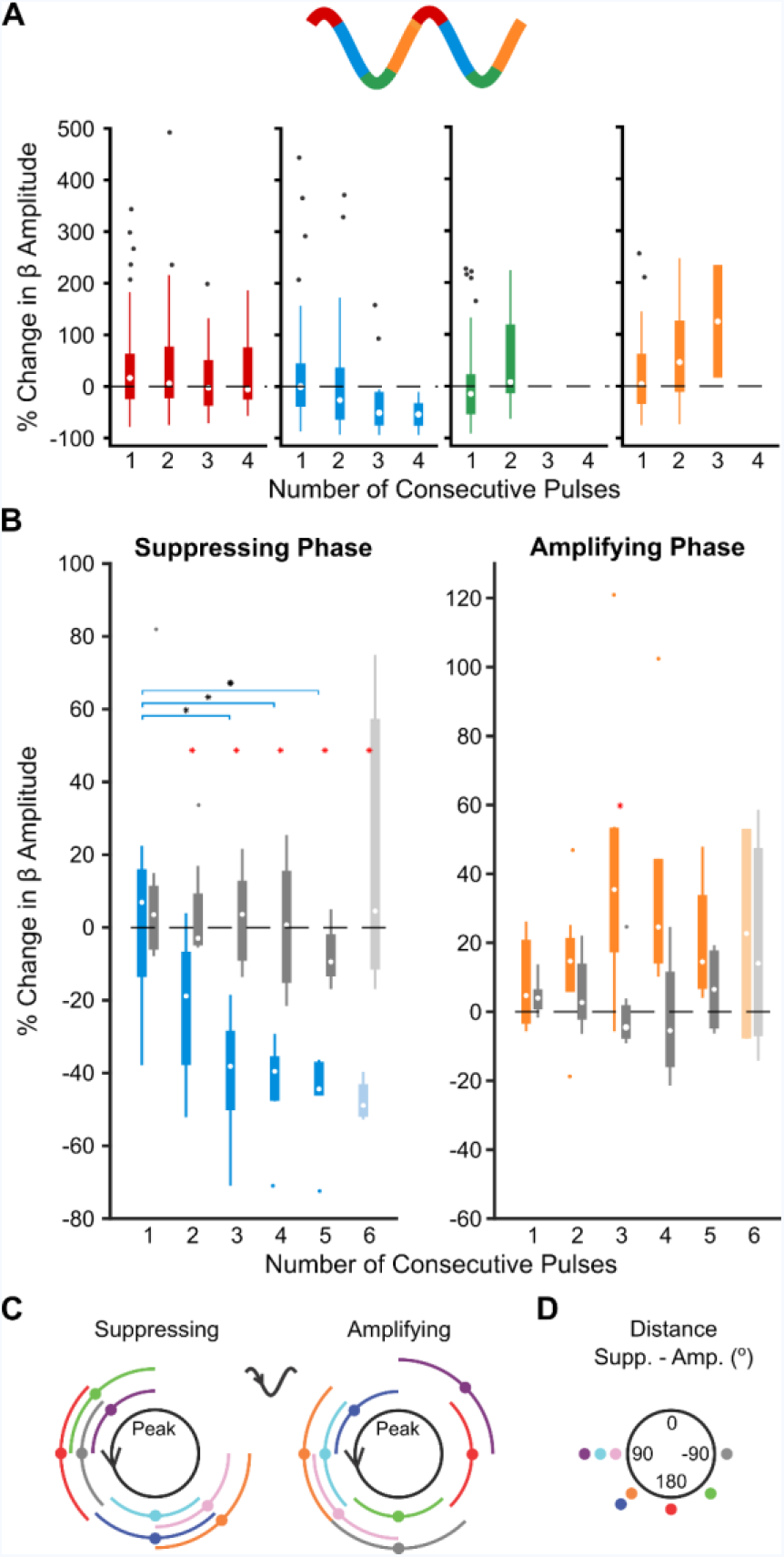
Consecutive phase-consistent stimulus pulses modulate beta oscillation amplitude. (**A**) Percentage change in beta oscillation amplitude from the median after consecutive cycles of stimuli occurring at a consistent phase is shown for four phase bins in an example subject. In this subject, amplitude suppression was seen after three consecutive cycles of stimuli delivered on the descending phase of the oscillation (blue), while amplification was seen after three consecutive cycles of stimuli delivered on the ascending phase (yellow). Stimuli delivered at alternative phases (red, green) did not result in modulation of the beta amplitude. **(B)** Median suppressing (blue) and amplifying (yellow) effects were grouped across eight patients. The percentage change in beta amplitude was compared to surrogate effects generated by replacing the amplitude envelope with a segment from an unstimulated time-matched portion of data (gray). Beta suppression was dependent on the number of consecutive stimuli delivered at the suppressing phase of the oscillation (p = 0.0016, Kruskal-Wallis test), while beta amplification was not (p = 0.1902, Kruskal-Wallis test). As six consecutive stimuli were only observed in four out of the eight patients at the suppressing phase and two out of eight patients at the amplifying phase (indicated by lighter boxes), these was not included in the Kruskal-Wallis test. Horizontal lines indicate differences between groups (corrected for multiple comparisons using the false discovery rate, p ≤ 0.05). Red stars indicate stimulation effects significantly different from surrogates (p ≤ 0.05, Wilcoxon ranked sum test). **(C)** Suppressing and amplifying phase bins for each patient. **(D)** Phase difference between the amplifying and suppressing phase bin for each patient.

Across patients, phase dependent suppression of the beta oscillation was dependent on consecutive cycles of stimulation at the suppressing phase (p = 0.0016, Kruskal-Wallis test). The mean percentage reduction in beta oscillation magnitude went from 21.8% after two consecutive stimulus pulses to 46.8% after five consecutive pulses. As it becomes increasingly unlikely that consecutive stimuli will arrive in the same phase bin, only four patients had at least 5 occurrences of 6 consecutive stimuli at the suppressing phase. However, in these four patients, 6 consecutive stimuli led to an average 47.5% reduction in amplitude. Importantly, beta amplitude reduction seen after two through six consecutive stimulus pulses at the suppressing phase was significantly beyond what was seen using the amplitude envelope from an unstimulated portion of the recording (p ≤ 0.05, Wilcoxon rank-sum test). This demonstrates that suppression was not simply the result of measuring amplitude after a certain number of consecutive cycles. Furthermore, a significant suppressive trend was seen at an individual level in five out of eight patients (p < 0.05, Kruskal-Wallis test) during stimulation at the suppressing phase, but in no patients when using the unstimulated amplitude envelope, p > 0.05, Kruskal-Wallis test). The strength of suppression across patients was inversely correlated with the relative spectral power at beta frequencies calculated across the entire recordings (r^2^ = 0.504, p = 0.049, linear regression analysis, Supplemental Fig. 6). This suggests that while suppression was still seen in patients with high spectral beta power overall, the instantaneous beta amplitude was more difficult to suppress than in patients with low relative peak beta power.

In contrast to suppressive effects, there was no significant effect of consecutive stimuli delivered at the amplifying phase (p = 0.190, Kruskal-Wallis test) (Fig. 5B). Additionally, amplification was only greater than surrogate effects after three cycles of stimulation at the amplifying phase (p = 0.0093, Wilcoxon ranked-sum test). Following the third consecutive cycle, amplification returned to surrogate levels, and only two out of eight patients showed amplification dependence on consecutive stimulus cycles (p < 0.05, Kruskal-Wallis). The strength of amplification across all patients was inversely correlated with the spectral beta power of the entire recording (r^2^ = 0.540, p = 0.038, linear regression analysis) (Supplemental Fig. 6). Together this indicates it was more difficult to further amplify the exaggerated beta signal.

It has been suggested that phase dependent modulation of neuronal oscillations may rely on stimulation induced changes to the phase of the oscillation (Azodi-Avval & Gharabaghi, 2015; Holt et al., 2016; D. Wilson & Moehlis, 2014). To address whether this mechanism could apply here, we investigated whether more phase slips, indicating discontinuities in the oscillation phase, were seen in the 15 ms following consecutive cycles of stimulation at the suppressing or amplifying phase (reported as a percentage of stimulus pulses) (Fig. 6A, B). In line with amplitude effects, there was no difference in the percentage of phase slips following the first stimulus (Fig. 6C, p = 1, Wilcoxon Ranked Sum Test). However, after the second and third consecutive pulse, significantly more phase slips are seen following stimuli arriving at the suppressing phase compared to the amplifying phase (Fig 6C, p = 0.0042, p = 0.0120, Wilcoxon Ranked Sum Test). In fact, almost no phase slips are seen following the third pulse at the amplifying phase, suggesting the oscillation is stable. The number of phase slips after consecutive cycles of suppressing stimulation was significantly different from surrogates (p = 0.0030, p = 0.0176, Wilcoxon Ranked Sum Test). Furthermore, after the second and third stimulus at the suppressing phase, the percentage of phase slips correlates with the reduction in beta amplitude (Fig. 6D, R^2^ = 0.0071, p = 0.0071; R^2^ = 0.61, p = 0.023; linear correlation), but not after the first pulse (R^2^ = 0.0018, p = 0.92) (Fig. 6D). At the amplifying phase, there was a significant inverse correlation with beta amplification, but only following the second stimulus pulse (Fig. 6E, R^2^ = 0.58, p = 0.028, linear correlation). These results are consistent with stimulation at the suppressing phase advancing or delaying the neuronal oscillation.

**Figure 6.**
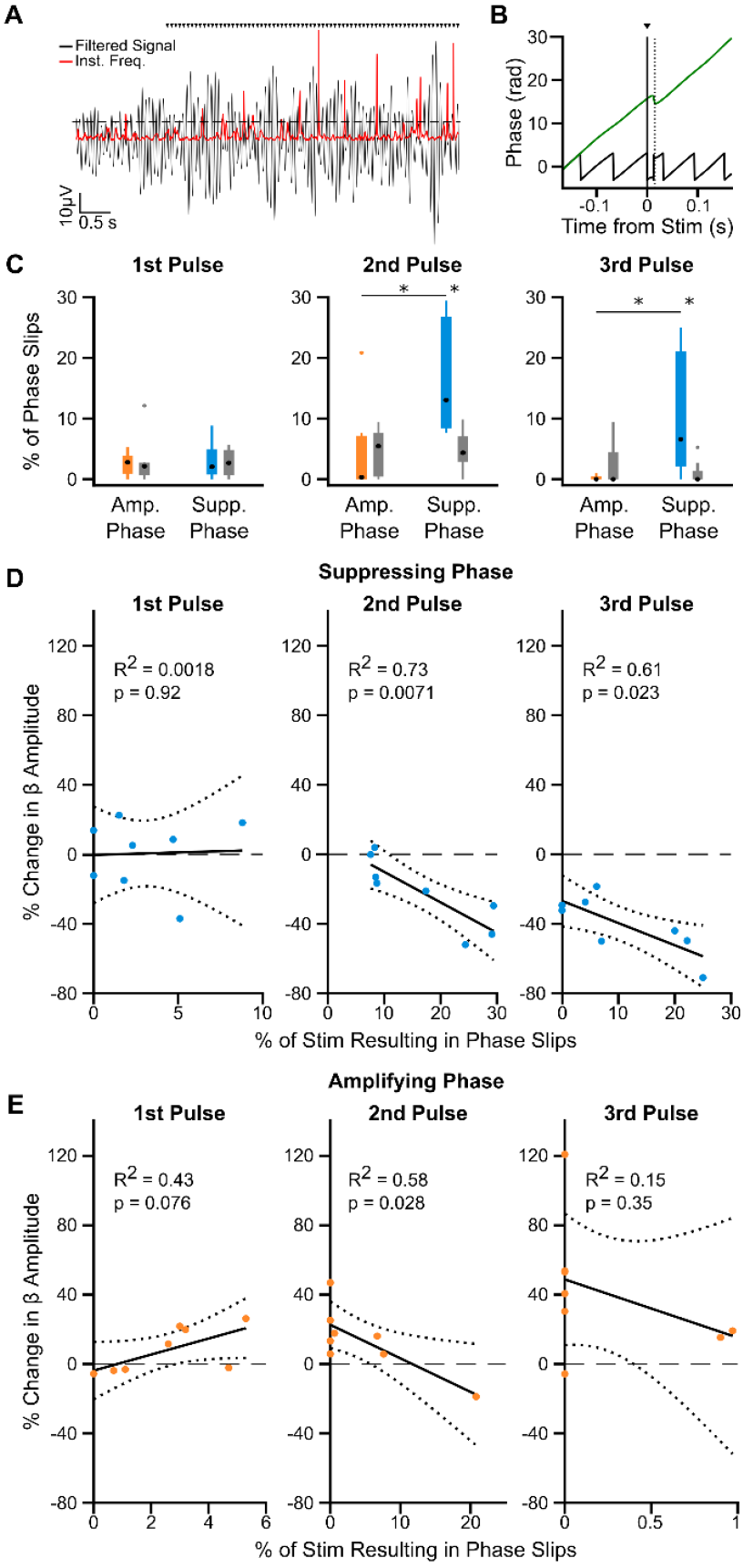
Increased phase slips in the beta oscillation following consecutive pulses at the suppressing phase correlates with amplitude reduction. (**A**) Phase slips were defined when the instantaneous frequency (red) of the beta filtered signal (black) crossed two standard deviations above the mean (dotted line), indicating a phase discontinuity in the oscillatory signal. Black triangles indicate stimulation pulses. **(B)** Example phase slip within 15 ms of a stimulus pulse (dotted line), seen in both the unwrapped phase (green) and phase (black) of the oscillation. **(C)** The percentage of stimulus pulses with phase slips occurring within 15 ms following the first (left), second (middle), and third (right) consecutive pulse at the amplifying (orange) or suppressing (blue) phase. Significantly more phase slips are seen after two and three pulses at the suppressing phase than at the amplifying phase (p = 0.0042, p = 0.0120 respectively, Wilcoxon Ranked Sum Test), and when compared to surrogates generated by replacing the signal with a time-matched unstimulated epoch from the same recording (p = 0.003, p = 0.0176, Wilcoxon Ranked Sum Test). **(D)** The percentage of phase slips occurring after the second (middle) and third (right) pulse at the suppressing phase correlates with the reduction in beta amplitude (p = 0.0071, p = 0.023, Linear Regression), but not following the first pulse (left, p = 0.92, Linear Regression). Note maximum x-axis values are different for each stimulus pulse as more slips occur following the second and third pulses. **(E)** The percentage of phase slips only correlates with beta oscillation amplification after the second consecutive pulse at the amplifying phase (middle p = 0.028, Linear Regression), not after the first (left) or third (right). Note the change in x-axis values as compared to those in D, as less phase slips occur at the amplifying phase.

### Increased phase-specificity enhances cumulative phase-dependent suppression of beta amplitude

Next, we investigated how the phase precision of stimulus pulses affects modulation of the LFP beta amplitude. To achieve this, we reanalyzed the suppression and amplification effects using stimulus-phase bins of different widths. Stronger effects using narrower bins would indicate dependence on the precision of the stimulus phase. In an example patient (Case 3) both enhanced suppression and amplification of the beta oscillation were seen when stimulus pulses occurred in a narrower window around the suppressing and amplifying phase (phase bins ⅛ th a cycle wide, amplifying bin: p = 0.0251, suppressing bin: p = 0.0272, Kruskal-Wallis test) (Supplemental Fig. 7A). Alternatively, when stimulus pulses occurred over a wider window around the suppressing and amplifying phase (phase bins ½ a cycle wide), neither significant amplification (p = 0.9413, Kruskal-Wallis test) nor suppression (p = 0.3882, Kruskal-Wallis test) was seen.

In order to observe cumulative effects of stimulation phase using phase bins ⅛ th of a cycle wide, the stimulus frequency had to be within 1 Hz of a patient’s peak beta frequency. As this was not always the case, it was not possible to provide group data using these narrower phase bins. However, bins could be widened to half a cycle for all patients (instead of a quarter of a cycle wide as in Fig. 5B). When stimulus pulses were allowed to occur over a wider window around the mean suppressing and amplifying phase, neither significant suppression nor amplification was seen across patients, even after consecutive pulses (Suppressing phase: p = 0.2755, Amplifying phase: p = 0.2502, Kruskal-Wallis test) (Supplemental Fig. 7B). Overall, these results suggest that at least a quarter cycle precision is needed to see phase dependent effects of stimulation, and that narrowing the phase window for stimulation further could lead to larger effects on amplitude.

### Phase-dependent suppression of beta amplitude was dependent on stimulation amplitude

To test whether phase-dependent suppression of beta oscillations is dependent on stimulation amplitude, three stimulus amplitudes (0.5 mA, 1 mA, and 1.5 or 2 mA) were applied while maintaining a consistent microelectrode location in three patients. As we found that suppression levels are affected by the strength of the beta power (Supplemental Fig. 7), it is important to view the differences across amplitudes within each patient instead of absolute values. Consistently across patients, stronger suppression was seen after the first, second, and third pulses occurred at the suppressing phase using 1 mA stimulation compared to 0.5 mA (Supplemental Fig. 8). In two of the patients (black and red), suppression seemed to plateau for each stimulation amplitude used. While in some cases, further suppression could be seen using 2 mA stimulation, results were not as consistent.

### Phase dependent suppression of oscillatory STN output

We next investigated whether a simultaneous change in unit activity could be seen concurrent with amplification and suppression of the LFP beta oscillation. For recordings included in the LFP analysis it was necessary to prioritize a recording location with strong beta oscillations, however, it was not always possible to maintain a stable single- or mutli- unit for the entire stimulation epoch. Therefore, background unit activity (BUA) was used as a measure of neuronal activity, as this could be used across all patients and represents output from a local population of neurons surrounding the recording contact. When stimulus pulses occurred at the suppressing phase of the LFP over consecutive cycles, the amplitude of beta oscillatory activity in the BUA decreased across patients (Fig. 7, p = 0.0046, Kruskal-Wallis test). However, instead of seeing suppressive effects by the second or third cycle, a decrease in BUA oscillatory activity was not seen until the fourth consecutive cycle, where a median 18.7% amplitude reduction was seen. Conversely, when stimulus pulses occurred at the amplifying phase of the LFP, enhancement of oscillatory activity in the BUA signal was not seen over consecutive cycles (Fig. 7, p = 0.7993, Kruskal-Wallis test). These results indicate that stimulating at the suppressing phase of the LFP results in a concurrent decrease in synchronous beta frequency output of STN neurons, and could therefore modulate oscillations in downstream structures and the wider network.

**Figure 7.**
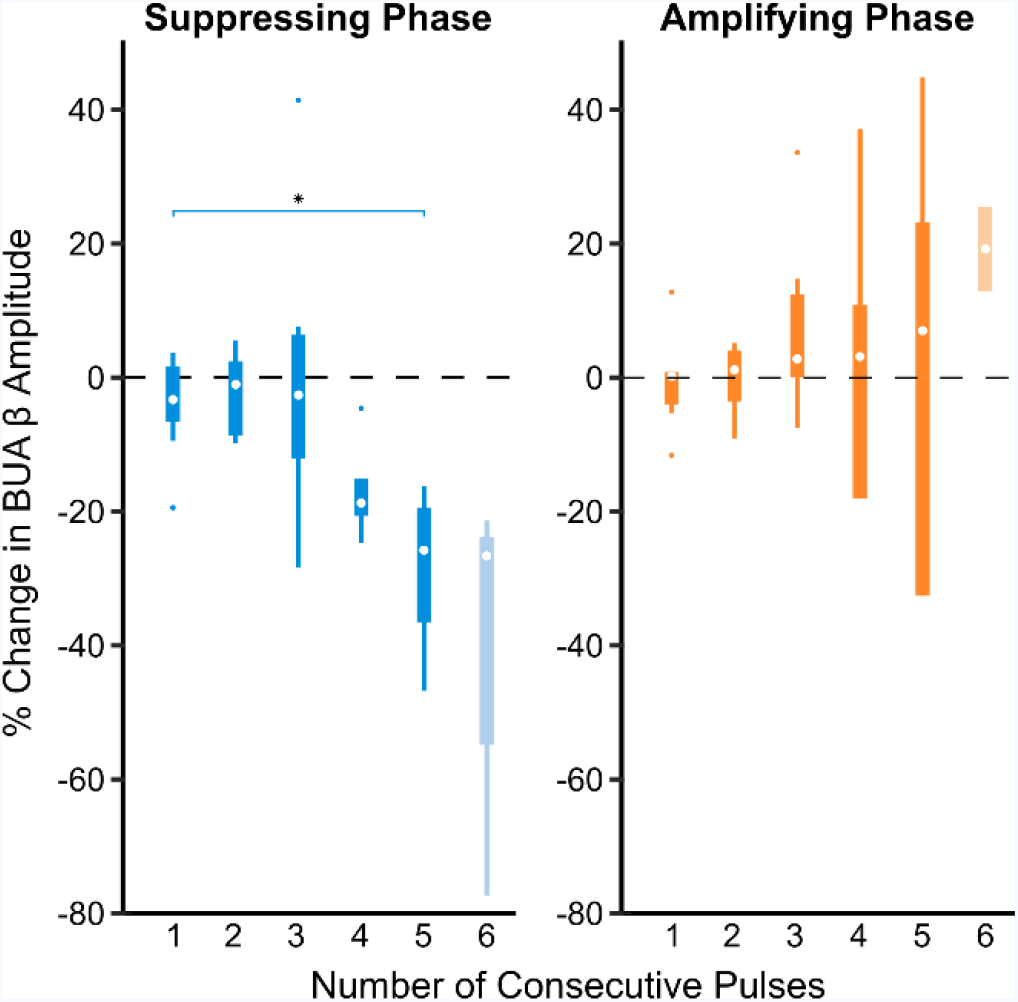
Consecutive stimulus pulses at the suppressing phase leads to suppression of the beta synchronous STN unit activity. (**A**) Background unit activity (BUA) was filtered around the peak beta frequency detected in the LFP. Median percent changes in the oscillation amplitude (compared to the median) following stimulation at the suppressing (blue) and amplifying (yellow) phase were grouped across eight patients. Suppression of beta frequency activity in the BUA was dependent on the number of consecutive stimuli delivered at the suppressing phase of the LFP oscillation (p = 0.0046, Kruskal-Wallis test), while amplification was not (p = 0.7993, Kruskal-Wallis test). As six consecutive stimuli were only observed in four out of the eight patients at the suppressing phase and two out of eight patients at the amplifying phase (indicated by lighter boxes), these was not included in the Kruskal-Wallis test. Horizontal lines indicate differences between groups (corrected for multiple comparisons using the false discovery rate, p ≤ 0.05).

### Phase dependent suppression led to decreased cortico-subthalamic beta-synchronization

Several studies have demonstrated that beta oscillations in STN are driven by those in the motor/premotor cortex (Fogelson et al., 2006; Shimamoto et al., 2013; Williams et al., 2002) and this synchronization is correlated with the severity of akinetic/rigid symptoms (Sharott et al., 2018). Using coincident EEG recordings, we examined whether changes in local oscillation amplitude induced by phase-dependent stimulation extended to effects on cortico-subthalamic synchronization. One patient only had Fz EEG recordings, so was not included in Pz and Cz analysis. In line with previous work (Ashby et al., 2001; Eusebio et al., 2009; Kumaravelu, Oza, Behrend, & Grill, 2018; Walker et al., 2012), stimulation pulses led to evoked potentials in midline EEG channels (Fig. 8A-C). While this demonstrates cortical activity could be modulated using our stimulation protocol, consecutive stimuli occurring at the suppressing or amplifying phase of the LFP beta oscillation were not associated with simultaneous changes in cortical beta amplitude (Fz: amplifying: p = 0.1269, suppressing p = 0.9526; Pz: amplifying: p = 0.7746, suppressing p = 0.9049; Cz: amplifying: p = 0.6275, suppressing: p = 0.9996; Kruskal-Wallis test). However, the phase alignment between midline EEGs and the STN during stimulation at the suppressing phase was less consistent during three cycles of stimulation at the suppressing phase than at the amplifying phase (Fig. 8D-F). This suggests phase dependent stimulation may have network-level effects on cortico-subthalamic synchrony in addition to local effects on STN beta oscillations.

**Figure 8.**
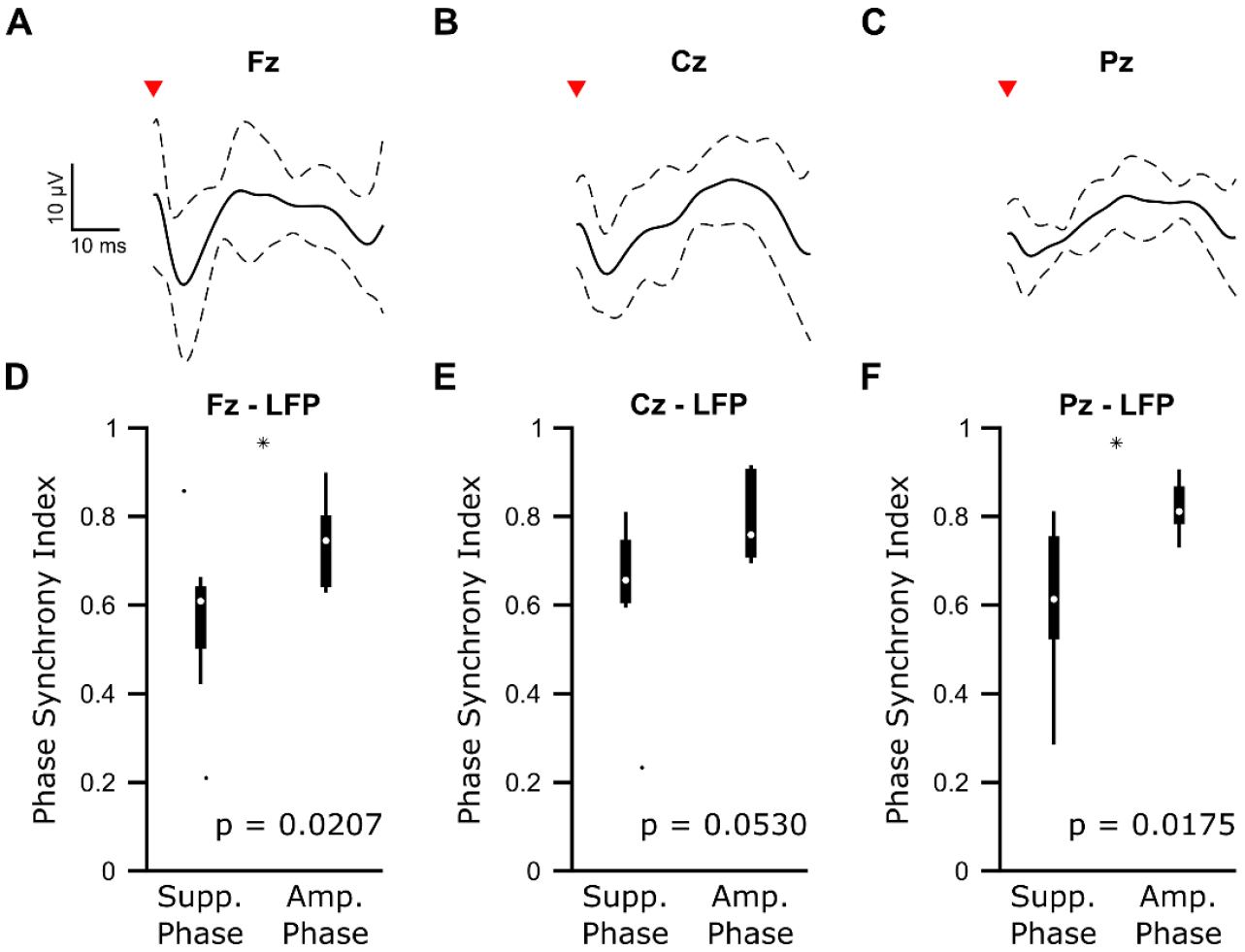
Phase dependent suppression of subthalamic beta oscillations is associated with lower cortico-subthalamic beta synchronization than phase dependent amplification. (**A-C**) Average (± standard deviation) cortical evoked responses from frontal/midline EEG (Fz, Cz, and Pz) locations) across eight patients for Fz and seven patients for Cz and Pz. EEG signals were bandpass filtered 1 – 100 Hz to remove any slow drift, and notch filtered 49 – 51 Hz to remove line noise. **(D-F)** Median phase synchrony index between the STN LFP and frontal/midline EEG channels over three cycles of consecutive stimuli occurring at the suppressing and amplifying phases across eight patients (Fz) or seven patients (Cz, Pz). Stimulating at the suppressing phase resulted in significantly lower cortico-subthalamic phase synchrony for Fz-LFP (p = 0.0207, Wilcoxon ranked sum test) and Pz-LFP (p = 0.0175, Wilcoxon ranked sum test, p < 0.05 indicated by black stars).

## Discussion

In this study we show that, in patients with Parkinson’s disease, electrical stimulation delivered to a specific phase of the subthalamic beta oscillation results in suppression of local and network level pathophysiological activity. Although beta frequency stimulation did not alter the power of beta oscillations over the entire recording, transient amplification and reduction of the oscillation amplitude could be seen when the specific timing of stimulus pulses was taken into account. Remarkably, delivering stimulus pulses locked to the suppressing phase over multiple consecutive cycles reduced the beta amplitude by over 40% across patients. This provides the first evidence that a phase-dependent approach to DBS could be more efficient and effective than high-frequency stimulation at reducing pathological neural oscillations in Parkinson’s disease. Moreover, our findings show that phase-dependent stimulation has the wider potential to disrupt or amplify neuronal oscillations to manipulate communication across brain areas.

### Potential mechanisms of stimulation

Any electrical stimulation of the brain, including high-frequency DBS, affects multiple neuronal elements in the vicinity of the stimulating electrode (McIntyre, Mori, Sherman, Thakor, & Vitek, 2004). While the novel stimulation set-up used in this study better allowed us to recover and analyze the underlying beta oscillation, the electrical current spread was likely different from conventional DBS. Due to the dorsal stimulating position, together with the horizontal orientation of the electric field (as opposed to the vertical orientation with conventional DBS electrodes), current was potentially delivered to multiple neuronal populations and fiber tracts (internal capsule, zona incerta, and fields of Forel) containing excitatory cortico-subthalamic and inhibitory pallido-subthalamic axons (Hamani, Saint-Cyr, Fraser, Kaplitt, & Lozano, 2004). Stimulation of these elements could lead to both orthodromic effects in the STN (and other targets) and antidromic effects at the source of the afferent fibers. Current may also have spread to the STN itself, but given the bipolar configuration it was likely more concentrated in these dorsal areas. Thus, as with therapeutic DBS, modulation of STN activity in our configuration likely occurred through a variety of direct and indirect mechanisms that cannot be fully delineated.

This combination of stimulation effects could potentially explain the variance in the multiphasic responses that our stimulation evoked in STN spiking activity, including both short latency (<10ms) excitation and inhibition. Such multiphasic responses result from a combination of excitatory and inhibitory afferents and the integration of these inputs with the pacemaker currents that drive the much of the spontaneous firing of STN neurons (Magill et al., 2004; Nambu et al., 2000). Overall, while neuronal activity in the STN was clearly influenced by the stimulation, it did not produce a consistent modulation in a particular direction of magnitude or frequency.

### Phase dependent amplitude suppression relies on consecutive pulses

Here we identify the existence of a critical phase of the STN beta oscillation, specific to each patient, that when stimuli occurred for multiple consecutive cycles leads to amplitude suppression of beta oscillatory activity in the LFP and BUA and a decrease in beta phase synchrony with the cortex. In parkinsonian patients and animal models of Parkinson’s disease, single STN neurons lock to a specific phase of the ongoing beta oscillation in the LFP (Kuhn et al., 2005; Mallet, Pogosyan, Sharott, et al., 2008; Sharott et al., 2018; Sharott et al., 2009; Weinberger et al., 2006), which leads to a correlation in the spiking across STN neurons at zero lag (Levy et al., 2002b; Mallet, Pogosyan, Marton, et al., 2008; Mallet, Pogosyan, Sharott, et al., 2008; Weinberger et al., 2006). The LFP likely reflects synchronized membrane currents of a population of neurons in the vicinity of the recording electrode. Several studies have demonstrated that oscillations in the motor cortices precede those in the STN LFP (Fogelson *et al.*, 2006; Lalo *et al.*, 2008; Litvak *et al.*, 2010; Sharott *et al.*, 2018), suggesting that cortical oscillations, measured here using EEG, may drive beta in the STN LFP through monosynaptic and or polysynaptic pathways. In contrast, the BUA must predominantly reflect spiking activity due to exclusion of low (>300Hz) frequencies (Moran et al., 2008; Siegel, Buschman, & Miller, 2015). In the context of STN beta oscillations, the EEG and STN LFP signals are therefore more indicative of synaptic input to the neurons, while the BUA signal reflects the output. In line with this delineation, previous work shows the BUA output can be vastly different in conditions with similar EEG/LFP power and synchrony, demonstrating that these signals are independent (Sharott et al., 2017).

Taken together, our results show that stimulation at consecutive pulses of the suppressive phase can reduce the oscillatory input to STN (STN LFP amplitude and cortico-subthalamic synchronization) and the output (BUA amplitude). Stimulus pulses delivered to neural oscillators can result in advances or delays in the oscillator depending on the stimulus phase (Ermentrout & Chow, 2002; Farries & Wilson, 2012; Smeal, Ermentrout, & White, 2010). In line with this theory, stimulation at the suppressing phase of the beta oscillation resulted in more disruptions to the phase following the second and third pulse. While this may be due to the fact that the oscillation is becoming less stable with amplitude reduction, both amplitude and phase slip effects were seen following the second stimulus. This makes it difficult to delineate which arises first, but provides further evidence that the timing of neuronal activity is being altered along with the amplitude. Within the STN, a phase shift in the beta oscillatory input may alter or reflect the reliability of the recruitment of STN neurons to the cortical oscillation, measured here using the BUA beta amplitude. Studies of beta oscillations in experimental animals have demonstrated that the synchronization observed in the STN extends across the entire cortico-basal ganglia network (Brazhnik, McCoy, Novikov, Hatch, & Walters, 2016; Deffains, Legallet, & Apicella, 2010; Goldberg et al., 2002; Mallet, Pogosyan, Marton, et al., 2008; Mallet, Pogosyan, Sharott, et al., 2008; Sharott et al., 2017; Tachibana, Iwamuro, Kita, Takada, & Nambu, 2011). Advancing or delaying the subthalamic beta output could thus disrupt the temporal relationship between downstream brain regions, which could result in decoupling of the network as a whole.

Importantly, we only saw these effects when two or more pulses were delivered on the suppressive phase. The simplest explanation for this is that the stimulus amplitudes delivered were generally low amplitude and, indeed, stronger beta amplitude suppression with fewer pulses was seen using higher stimulation amplitudes. However, it is also possible that disrupting the temporal relationships between oscillators within and between nodes of the network may require several cycles to decouple the circuit. Regardless, one key aspect in developing improved stimulation protocols for DBS is reducing the stimulation current to prevent spread to regions outside the target. Our results suggest delivering low amplitude pulses at the suppressing phase over several cycles is sufficient to disrupt pathological network activity. Interestingly, unlike suppressing effects, phase dependent amplification did not depend on the number of consecutive cycles of stimulation and was generally weaker than suppression. This may be because oscillatory activity is already in a heightened state in Parkinson’s disease, making it more difficult to enhance than suppress. The linear relationship seen between beta power and amplification resulting from stimulation supports this hypothesis. This property is potentially therapeutically useful, as enhancing ongoing beta would potentially worsen symptoms.

### Implications for therapy

Advancements to DBS algorithms have been aimed at improving efficacy and reducing the amount of current delivered, which could limit stimulation-induced side effects and conserve battery power (Adamchic et al., 2014; Brocker et al., 2017; Cagnan et al., 2017; Little et al., 2013; D. Wilson & Moehlis, 2014). Closed-loop approaches to neurostimulation have already proven to be achieve some of these aims (Little et al., 2013; Rosin et al., 2011). For a closed-loop approach to be effective, an accurate biomarker of symptom severity must be used to drive stimulation. While a number of synchrony measures have been implicated in motor symptoms of Parkinson’s (de Hemptinne et al., 2013; Kühn et al., 2009; Tass, 2003; Weinberger et al., 2006), beta oscillations offer an appealing biomarker for the control of akinesia and rigidity for a number of reasons. First, beta oscillations can be recorded in the LFP and have proven to be an effective chronically recorded feedback signal in patients even during unimpeded movement (Little et al., 2013; Rosa et al., 2015). Second, recordings can be made from the implanted stimulation electrode, limiting the operation to a single surgical site. Finally, alternative measures implicated in motor symptoms, specifically coupling between the amplitude of high frequency activity to the phase of the beta oscillation (phase amplitude coupling) (de Hemptinne et al., 2013), would likely be disrupted with the suppression of the carrier frequency.

## Conclusions

Our findings suggest that tracking the phase of ongoing beta oscillations and delivering stimulus pulses only on the suppressing phase could be an effective closed-loop strategy in Parkinson’s disease. Implementing such phase tracking would allow a full evaluation of the efficacy of phase-dependent stimulation for selectively reducing beta and sparring functional activity. The increased number of consecutive stimulation cycles possible with active phase tracking could further enhance suppressive effects. Furthermore, unlike tremor, where there is an ongoing measure of motor impairment, akinetic/rigid symptoms require task-related evaluation. Phase-tracking would allow longer periods of phase-locked stimulation (compared to the 100 – 400 milliseconds seen in this study) that would enable this type of behavioral assessment. While active phase locked stimulation has been applied to low frequency neural oscillations in the hippocampus (Siegle & Wilson, 2014) and low frequency peripheral oscillations (Brittain et al., 2013; Cagnan et al., 2017), to our knowledge, this approach has not yet been successfully implemented at frequencies above 10 Hz in experimental animals or humans. Implementing active phase tracking of parkinsonian beta oscillations presents challenges not only due to the higher frequency signal, but also because oscillations tend to occur in short bursts (Tinkhauser et al., 2017) and signals are on the order of 1 μV. Such issues could be addressed by combining phase tracking with proven amplitude-based approaches to further optimize the selectivity of stimulation (Little et al., 2013). Overall, the results presented here provide strong evidence in support of exploring such an approach in the future, both for PD and other diseases where abnormal oscillations are a pathophysiological feature.

## Acknowledgements

The authors would like to thank all the patients who participated in this study.

---

**Supplemental Table 1.** Patient details.

| Case | Age (years), gender | Disease duration (years) | Motor UPDRS OFF | Motor UPDRS ON | Preoperative anti-parkinson drugs | Hoehn/Yahr Score | Dominant Side | Mattis Dementia Rating Scale | Major Symptoms | Included in analysis |
| --- | --- | --- | --- | --- | --- | --- | --- | --- | --- | --- |
| 1 | 72, M | 6 | 33 | 13 | Levodopa 650 g | 3 | Right | 143 | Akinetic, Rigid | Yes |
| 2 | 64, M | 7 | 50 | 19 | Levodopa 250 mg<br>Budipin<br>Piribedil<br>Rasagilin | 3 | Left | 127 | Akinetic, Rigid | Yes |
| 3 | 70, F | 9 | 30 | 12 | Levodopa 1350 mg | 3/4 | Left | 137 | Rigid, Tremor | Yes |
| 4 | 60, M | 12 | 31 | 17 | Levodopa 1000 mg | 2 | Right | 141 | Akinetic, Rigid Tremor | Yes |
| 5 | 68, F | 8 | 18 | 13 | Levodopa 500 mg | 3 | Right | 139 | Akinetic Rigid | Yes |
| 6 | 56, F | 10 | 34 | 16 | Levodopa 450 mg | 2 | Right | 139 | Akinetic Rigid | Yes |
| 7 | 57, M | 17 | 33 | 16 | Levodopa 1600 mg | 3 | Right | 143 | Akinetic, Rigid, Tremor | Yes |
| 8 | 62, F | 17 | 60 | 33 | Levodopa 1400 mg<br>Apomorphine<br>Pramipexol | 4 | Left | 137 | Akinetic, Rigid, Tremor | Yes |
| 9 | 53, M | 22 | 63 | 9 | Levodopa 600 mg<br>Benserazide 25 mg<br>Rasagiline 1 mg | 3 | Right | 137 | Akinetic, Rigid, Tremor | No |
| 10 | 49, M | 10 | 21 | 8 | Levodopa 600 mg<br>Benserazide 25 mg<br>Ropinirole 32 mg<br>Safinamide 100 mg | 2 | Right | 142 | Akinetic Rigid | No |

**Supplemental Figure 1.**
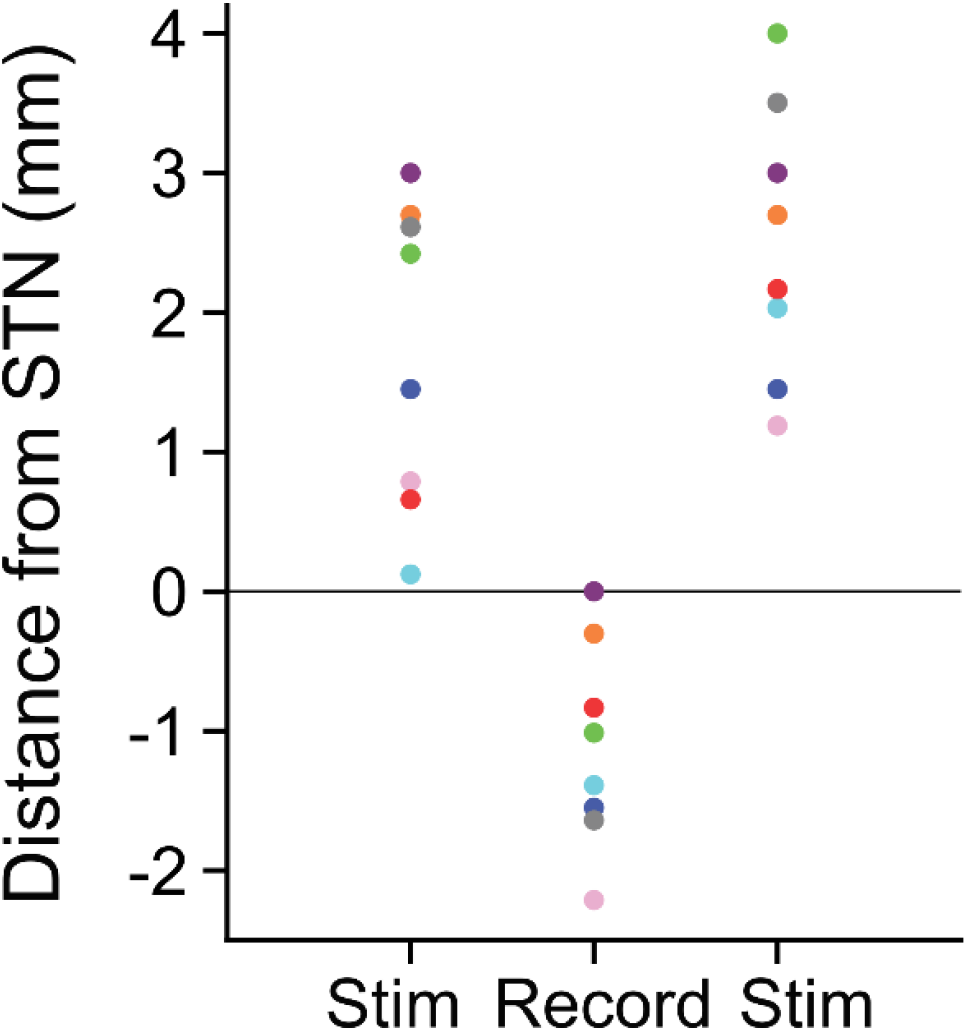
Bipolar stimulation was delivered dorsal to the subthalamic nucleus (STN) while recording within the STN. The dorsal STN border (set to zero) was defined for each patient based on physiological activity. For each patient, the distance of the two stimulation electrodes and recording electrode from the dorsal border is recorded. Patient colors are consistent with those used in Fig. 5C.

**Supplemental Figure 2.**
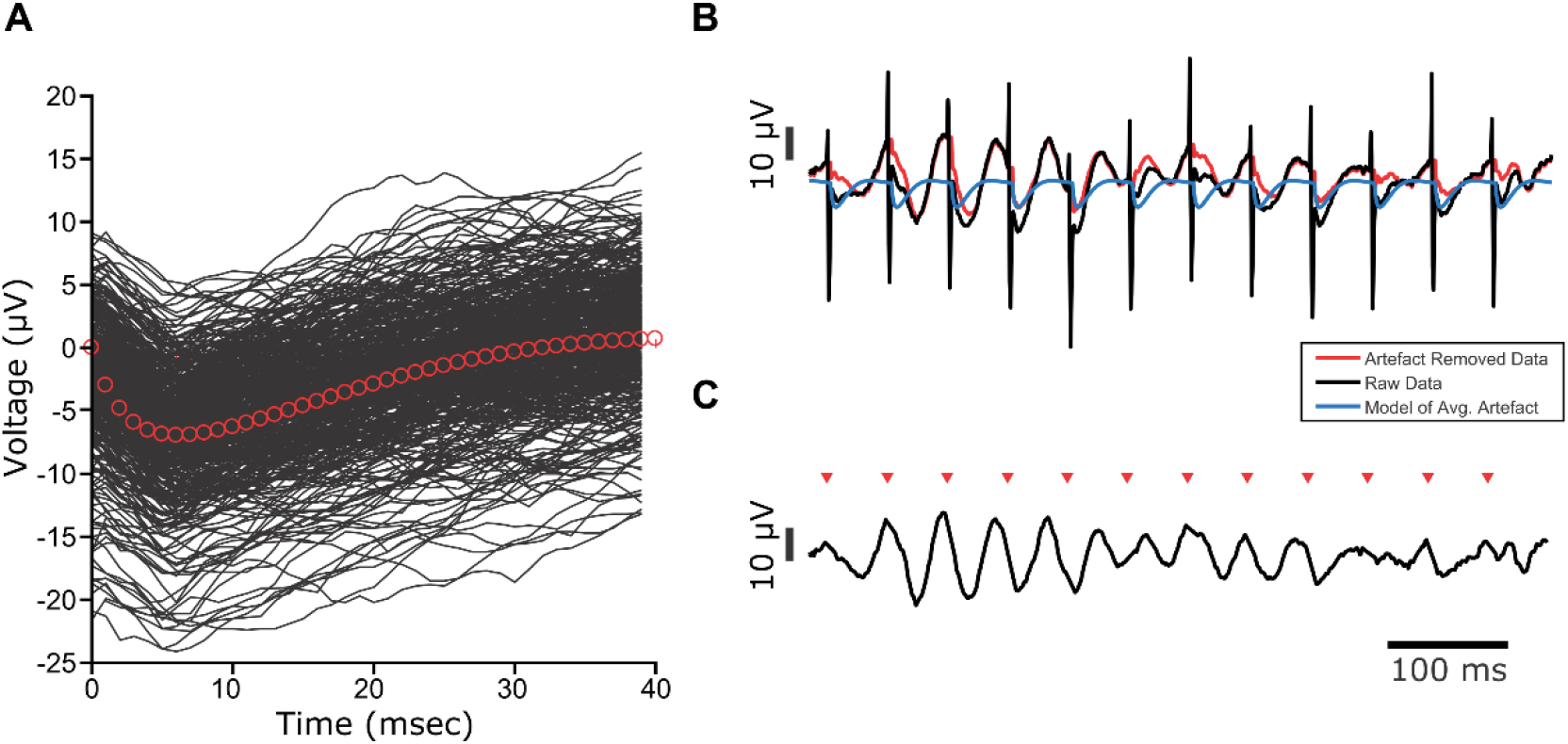
Stimulus evoked artifacts in the local field potential recordings were removed using a Kalman filter approach. (A) A transfer function model was fit to the average of all stimulus evoked artifacts. **(B)** A Kalman filter was used to generate an artifact free signal (red) using the raw signal (black) and a model of the average artifact (blue). **(C)** Voltage trace showing the artifact free signal (arrows indicate stimulus pulses).

**Supplemental Figure 3.**
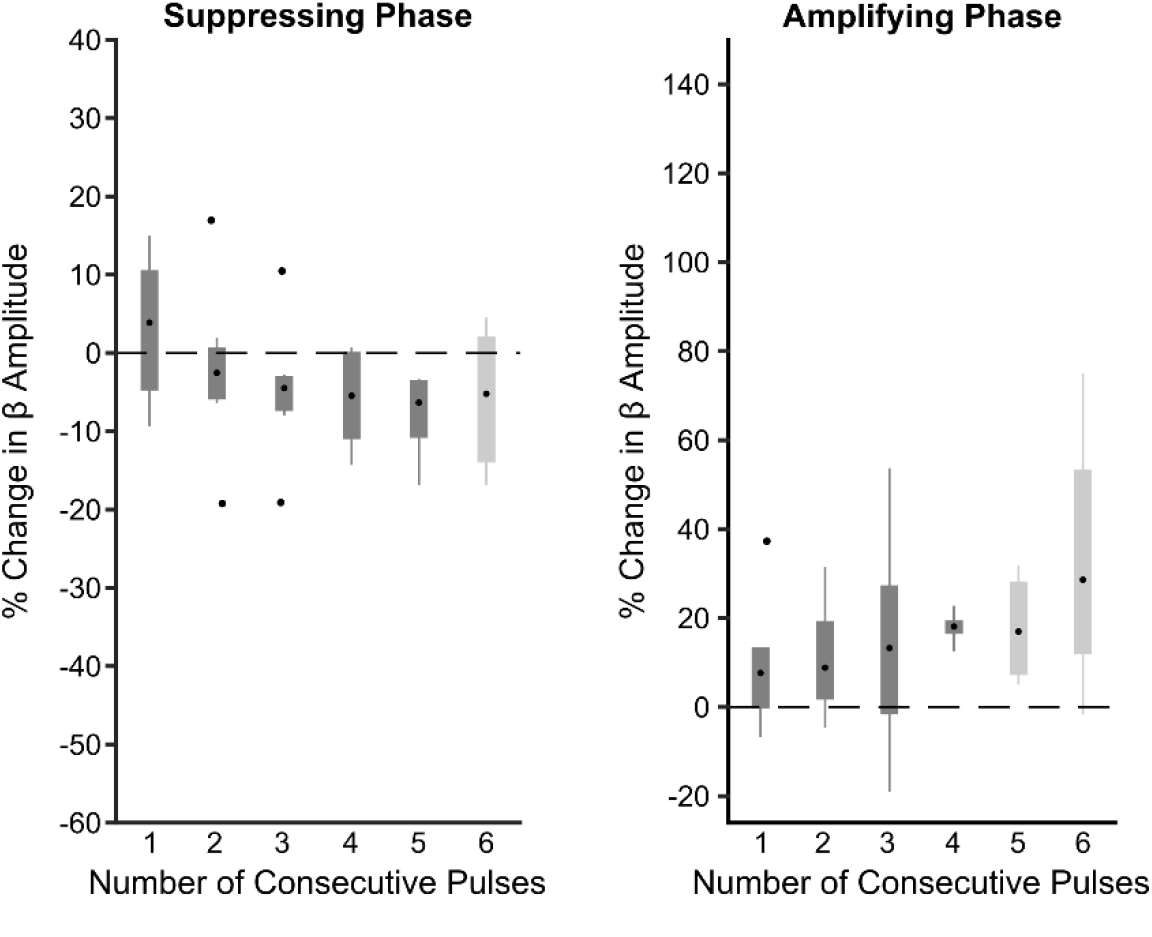
Significant beta amplitude modulation is not seen using alternative surrogates. The percent change in amplitude from the median is plotted as a function of number of consecutive stimulus pulses. The amplitude envelope from an unstimulated portion of the recording was used to replace the envelope during stimulation. Instead of preserving the amplification and suppression phases (as in Fig. 5), new suppression and amplification phases were chosen using the same methods described to select the phases during stimulation. While Fig. 5 allowed us to determine whether the same number of cumulative pulses lead to suppression or amplification because of the consecutive structure, this control allows us to assess the maximum phase dependent modulation of the beta amplitude seen without stimulation. Neither suppression (p = 0.3753) nor amplification (p = 0.6114) was dependent on the number of consecutive cycles of stimulation (Kruskal-Wallis test, p> 0.05). At the suppressing phase, six consecutive pulses were not seen in five patients, while at the amplifying phase neither five nor six consecutive pulses were seen in five patients (indicated by the lighter gray).

**Supplemental Figure 4.**
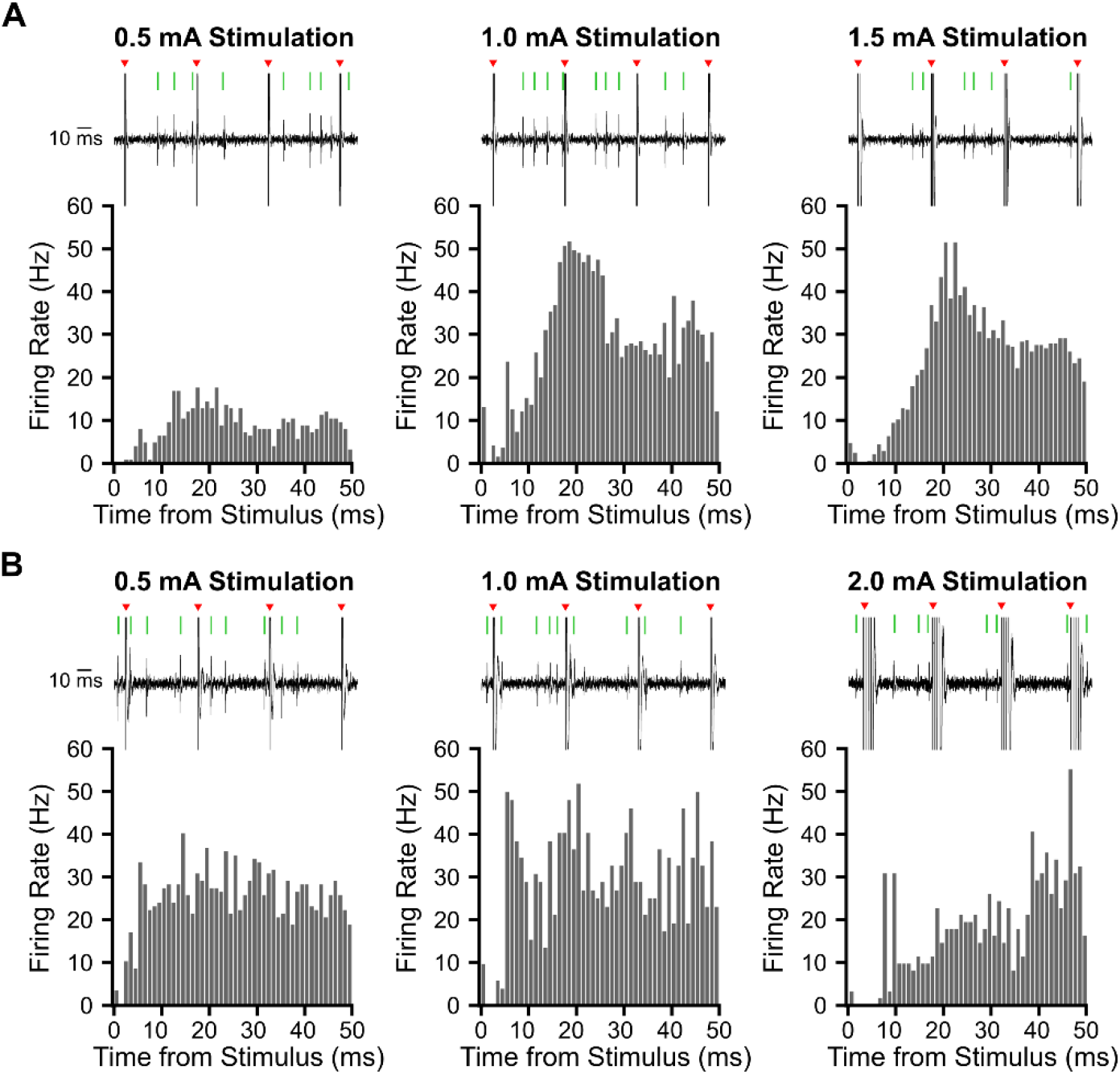
Modulation of subthalamic (STN) units by low amplitude beta frequency stimulation delivered dorsal to the STN can be amplitude dependent. Peristimulus time histograms (PSTH), using 1 ms wide bins, from two example units (2 patients) in response to three different stimulus amplitudes. Spikes were detected from microelectrode recordings in the STN; representative examples of raw unit data during 3 consecutive electrical stimuli are shown above each PSTH (black: raw trace; red arrow: stimulation; green line: detected spike). Beta frequency stimulation was delivered dorsal to the STN at **(A)** 0.5, 1, and 1.5 mA in subject 7. **(B)** 0.5, 1, and 2 mA in subject 1.

**Supplemental Figure 5.**
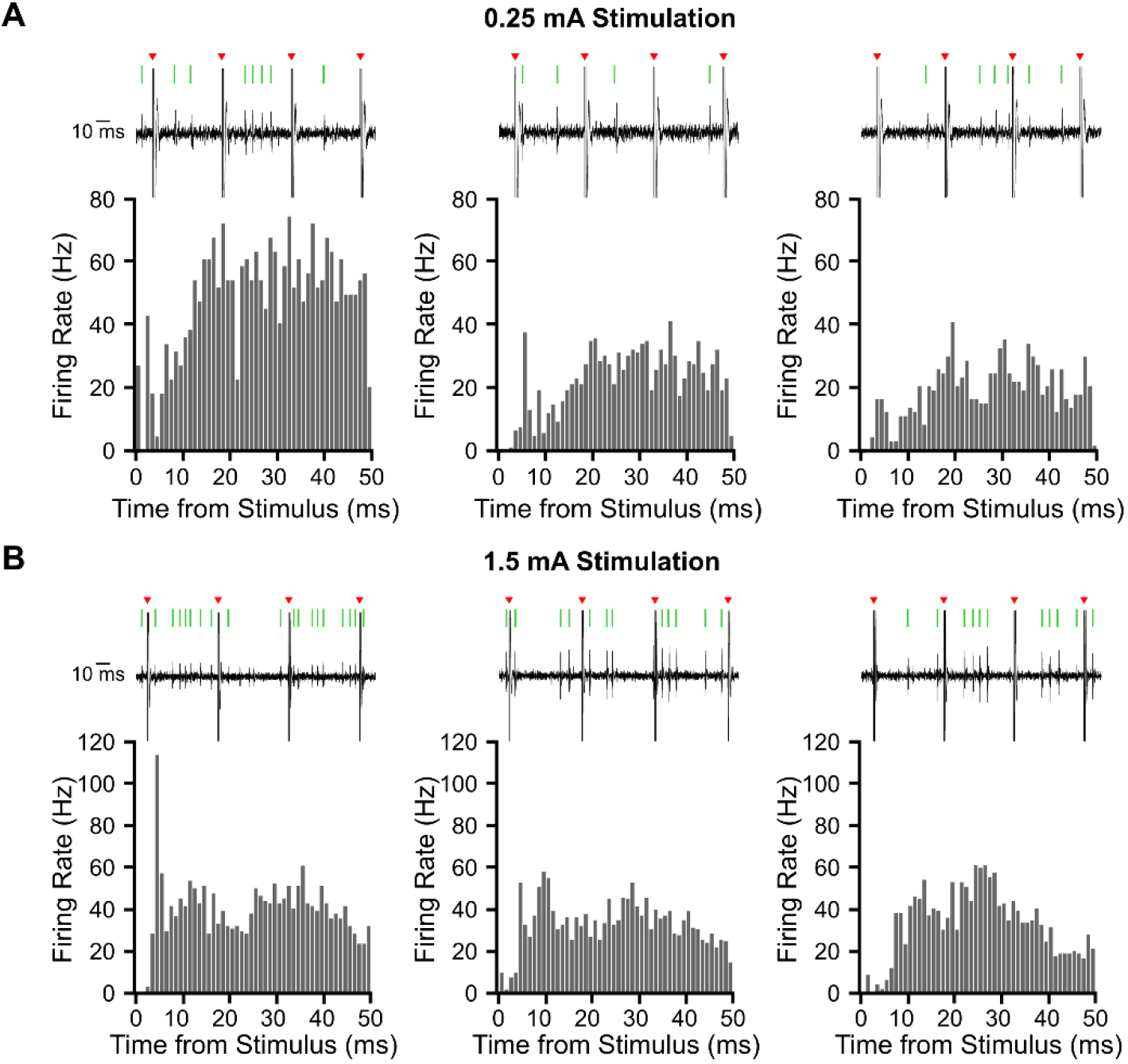
Modulation of subthalamic unit activity is inconsistent across different stimulation/recording locations within the same subject. In two patients, peristimulus time histograms (PSTH), 1 ms wide bins, are shown from three stimulation/recording locations (stimulation fixed at 3 mm above recording location). Spikes were detected from microelectrode recordings in the STN; representative examples of raw unit data during 3 consecutive electrical stimuli are shown above each PSTH (black: raw trace; red arrow: stimulation; green line: detected spike). Bipolar stimulation was delivered dorsal to the STN **(A)** at 0.25 mA for three STN units recorded from subject 4, **(B)** at 1.5 mA for three STN units recorded from subject 6.

**Supplemental Figure 6.**
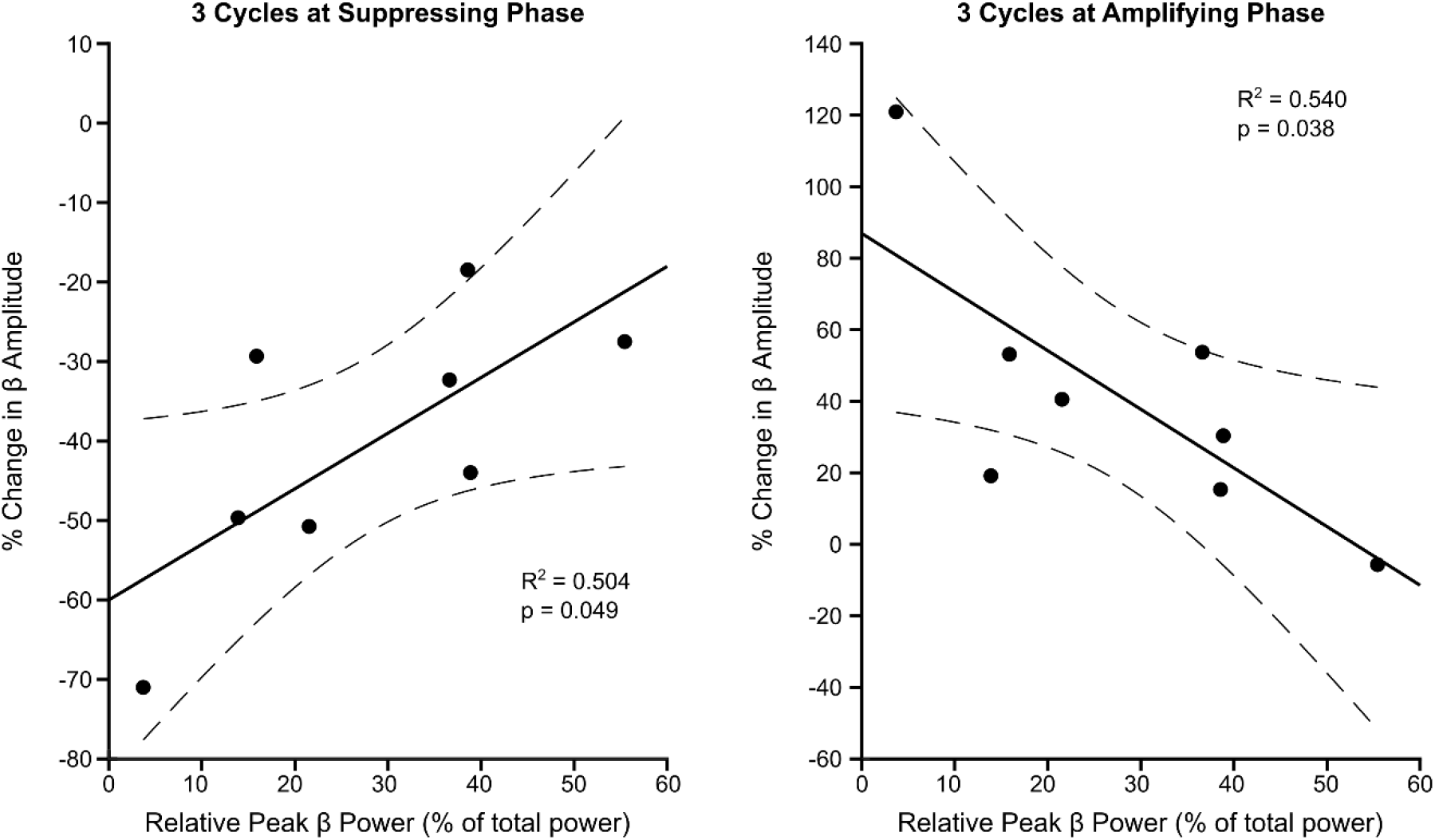
Correlation between the strength of beta amplitude modulation and the peak beta power. The percent change in beta amplitude after the third cycle of stimulation at the suppressing (left) and amplifying (right) phase is plotted as a function of the relative peak beta power (peak ± 3 Hz). The third stimulus pulse was chosen as this was the maximum suppression and amplification that could be seen in all patients (i.e. not all patients had 4 consecutive cycles of stimulation at the suppressing or amplifying phase). The peak beta power was calculated from the power spectrum of the entire recording (relative to 5 – 45 Hz), as seen in Fig. 2C. Linear regression fit is shown by the black line and the 95% confidence limits are shown with the dashed lines for the suppressing phase (R^2^ = 0.504, p = 0.049) and amplifying phase (R^2^ = 0.540, p = 0.038). This suggests stronger suppression is seen when beta power is lower, although suppression is still seen in the patients with high beta power. Furthermore, it is easier to amplify the beta amplitude of patients with less beta power, perhaps because it is difficult to further amplify the beta oscillation beyond a certain point.

**Supplemental Figure 7.**
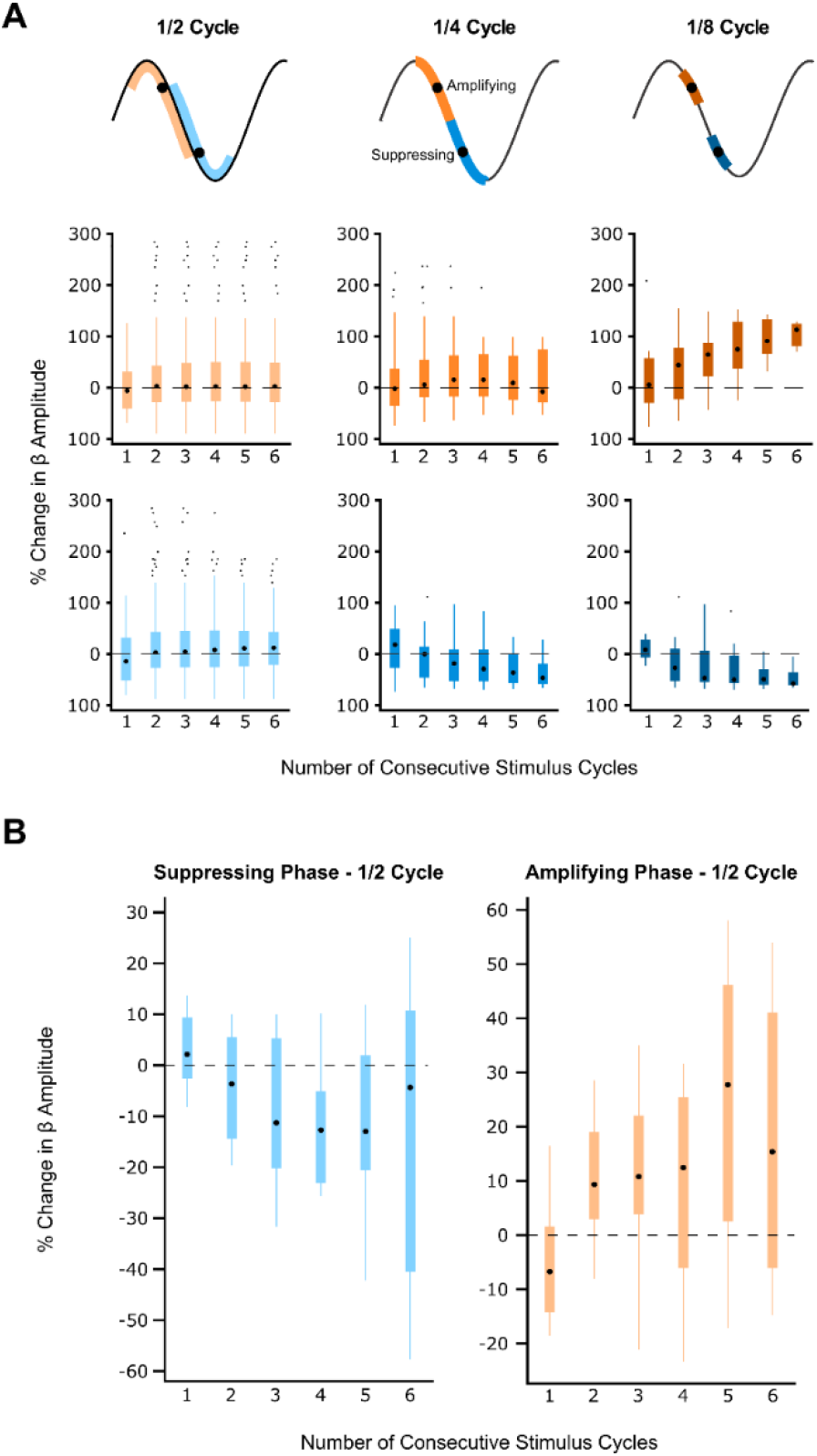
Phase precision of the stimulus pulse affects the strength of beta amplitude modulation. (A) Example patient showing both enhanced suppression and amplification of beta amplitude when using narrower phase bins. Stimulation frequency (20 Hz) was 1 Hz different from peak beta frequency (19 Hz), allowing for detection of consecutive pulses despite narrower phase bins. Bins were widened or narrowed around the mean suppressing and amplifying phase (black dots). Stimulus phase was defined as: left) half the oscillation cycle; middle) ¼^th^ the oscillation cycle; right) ⅛^th^ the oscillation cycle. Orange hues represent the suppressing phase; blue hues represent the amplifying phase. Amplitude modulation was only dependent on number of consecutive pulses when using ⅛^th^ of the oscillation cycle (amplifying phase: 2 bins, p = 0.4594; 4 bins, p = 0.972; 8 bins, p = 0.0251; suppressing phase, 2 bins, p = 0.8508; 4 bins, p = 0.0517; 8 bins, p = 0.0272, Kruskal-Wallis test) **(B)** Median suppressing and amplifying effects on the beta amplitude using phase bins half a cycle wide were grouped across eight patients. Neither beta suppression (p = 0.2755) nor amplification (p = 0.2502) was dependent on the number of consecutive stimuli (Kruskal-Wallis test). This is in contrast to results seen in Fig. 5 where narrower phase bins were used.

**Supplemental Figure 8.**
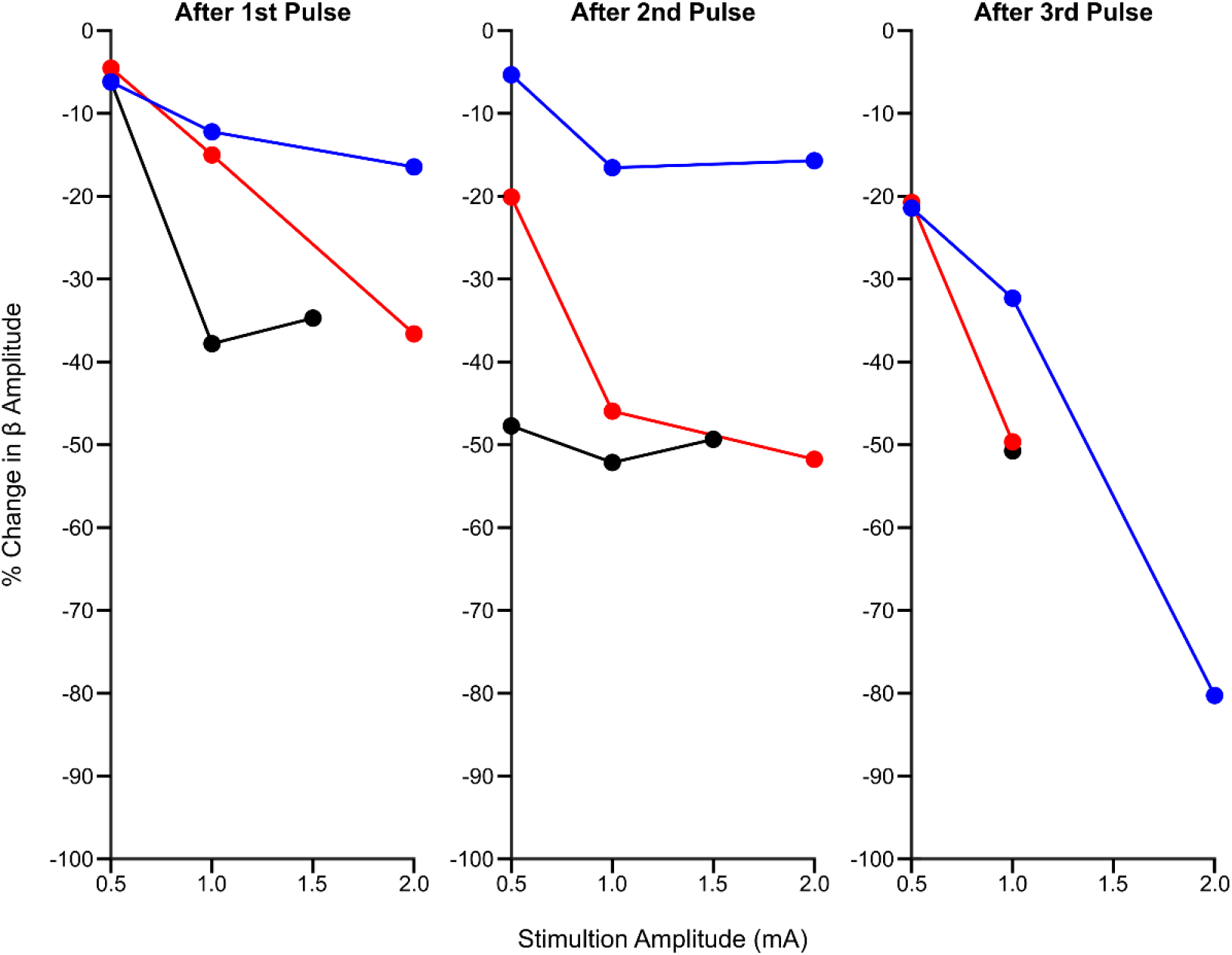
Phase dependent suppression of beta oscillations is dependent on stimulation amplitude. Percentage change in beta oscillation amplitude after the 1^st^ (left), 2^nd^ (middle), and 3^rd^ (right) consecutive stimulus pulse at the suppressing phase as a function of stimulus amplitude is plotted for three patients. After each stimulus, the decrease in beta amplitude was stronger when using 1 mA versus 0.5 mA. Further suppression could be seen when the stimulus amplitude was increased to 2 mA; however, effects were not as consistent.

## Abbreviations

STN: Subthalamic nucleus
DBS: Deep brain stimulation
LFP: Local field potential

